# On the transient interactions of α-synuclein in different dimensions

**DOI:** 10.1101/2025.01.29.635265

**Authors:** L. Ortigosa-Pascual, N. Ferrante-Carrante, K. Bernfur, K. Makasewicz, E. Sparr, S. Linse

**Affiliations:** Lund University

## Abstract

α-Synuclein (αSyn) is a neuronal protein predominantly found in the brain, whose native function seems to be associated with vesicle trafficking. While intrinsically disordered in solution, the first ca. 100 residues adopt an amphipathic α-helical structure when the protein adsorbs onto membranes. Additionally, the aggregation of αSyn into highly ordered β-sheet rich amyloid fibrils is associated with Parkinsońs disease. The different regions of αSyn and the interactions between them have been reported to play a key role in the behaviour of the protein in solution, its membrane binding, and its aggregation into fibrils.

This study employs photo-induced cross-linking of unmodified proteins (PICUP) to capture and identify the transient contacts of αSyn in different conformational states: free in solution, adsorbed to membranes, and aggregated into fibrils. By using tyrosine-to-phenylalanine mutations to block the reactivity of specific amino acid residues, we establish key cross-links in each state. In solution, we identify internal contacts between the N and C termini of monomers, as well as inter-monomer contacts between C termini in oligomers. When αSyn is adsorbed to membranes, the internal cross-linking is blocked, while cross-linking between C-terminal regions persists. In fibrils, cross-linking is significantly reduced, primarily occurring between C-terminal residues of adjacent monomers. This work highlights the utility of PICUP for reporting on the transient contacts that occur on the pathways of self- and co-assembly of αSyn.

## 1. Introduction

α-Synuclein (αSyn) is a neuronal protein associated with vesicle trafficking. Its aggregation has been linked with several neurodegenerative diseases, having been detected in inclusion bodies in the brains of Parkinson’s disease patients^1^, as well as the identification of a single point mutation of αSyn being linked to a case of familial Parkinson’s disease ^2^. αSyn has since been the target of many studies and has been thoroughly characterized in terms of membrane binding, aggregation and structure^3–9^.

αSyn is a 140 amino acid protein of the synuclein family with a highly asymmetric charge distribution (Figure 1C). The sequence can be divided into three regions based on its fibril formation propensity, consisting of an N-terminal tail (Nt), a hydrophobic fibril core, and a C-terminal tail (Ct). The Nt is net positively charged, with 10 lysine and 8 acidic residues. Its high conservation through the protein’s evolution suggests the Nt plays a critical role for the healthy function of αSyn and protects against disease^10^. This is supported by the fact that all familial Parkinson’s disease variants have a mutation in this region^2,11–16^. The central region, or fibril core, has a high content of hydrophobic residues, which play a central role in the stacking of monomers into β sheets to form amyloid fibrils. Residues 1-95 include seven repeats of a motif with the consensus sequence KTKEGV. These repeats have been shown to contribute to the lipid membrane binding of the protein both *in vivo*^17^ and *in vitro*^18^. The Ct, with 15 acidic amino acids, is strongly negatively charged at neutral pH and the pKa values are upshifted upon fibril formation^19^, which reduces the electrostatic repulsion in the fibril. The interaction between the oppositely charged N and C termini, be it within or between proteins, has been shown to play a key role in αSyn self-assembly^20–24^.

**Figure 1.**
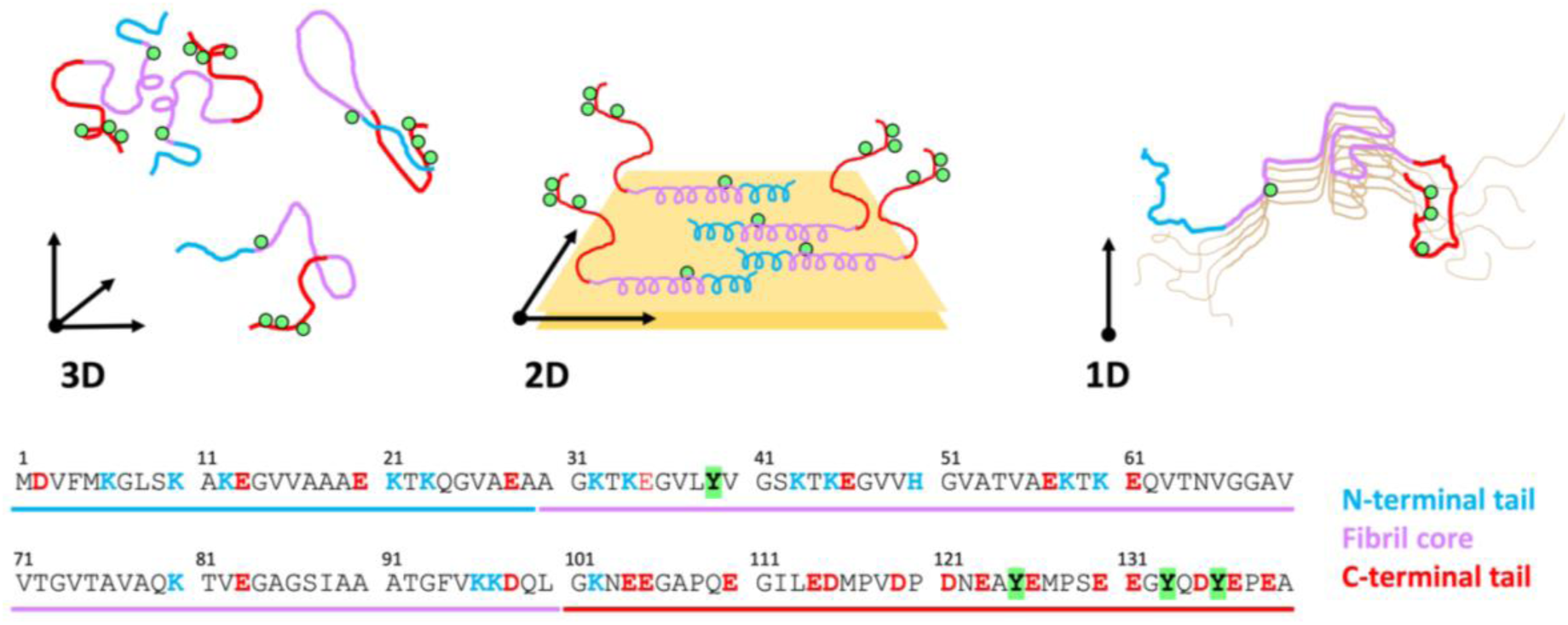
Cartoon representations of αSyn in solution (A), bound to membrane (B) and in fibril structure (C). The colour coding represents the N-terminal tail (blue), the fibril core (purple) and the C-terminal tail (red), with the tyrosine residues, targets of the PICUP reactions, depicted as green spheres. The proteins interact with each other in 3D (A), 2D (B) and 1D (C) depending on the system depicted. (D) αSyn sequence color coded to depict N-terminal tail (blue), fibril core (purple) and C-terminal tail (red). Negatively charged residues are shown in red, positively charged residues in blue, and tyrosines 39, 125, 133 and 136, main targets of the PICUP reaction, are highlighted in green.

Being an intrinsically disordered protein, monomeric αSyn doesn’t adopt a particular structure, but rather exchanges rapidly between different conformations^25^ (Figure 1A). This has been found to be the case also *in vivo*^26^. When αSyn adsorbs onto membranes, the first ca. 100 residues adopt an amphipathic α-helix structure, with the remaining amino acids in a disordered conformation^27^ (Figure 1B). This transition can be monitored with circular dichroism spectroscopy, showing a shift from the spectrum typical of a random coil to that of an α-helix.

The helix sits parallel to the membrane interface, with the center of the helix positioned slightly below the phosphate groups^3,4^. Furthermore, it was demonstrated that the association takes place when the membrane contains anionic lipids, while there is no adsorption to membranes containing only zwitterionic lipids^27^. The disordered Ct stick towards the solution forming a polymer brush of an estimated thickness of 6 nm^3^. The complexity of the αSyn-lipid interactions is considerable^9^. For instance, different αSyn binding modes have been observed, involving a different number of residues in the α-helix formation depending on the available lipid surface^28,29^. Recent work has also shown that αSyn binding to membranes occurs with positive cooperativity, meaning that the affinity of a protein is higher for a membrane with a previously bound αSyn molecule over a bare membrane^30^. Improving our understanding of the αSyn-lipid interactions is crucial to shed light into the αSyn activity in membrane remodeling and vesicle trafficking. The finding that Lewy Bodies, initially suspected to be mainly formed by αSyn, have a high lipid content accentuates this need^31^.

αSyn is an amyloid protein with propensity to aggregate into fibrils. Although different morphs of αSyn fibril structures exist ^24,32,33^, they all share the stacking of monomers forming extended β-sheets parallel to the fibril axis^32,34^ (Figure 1C). This aggregation is driven by hydrophobic interactions in the fibril core, while the negatively charged Ct remains disordered and extends into solution, forming a “fuzzy coat” with elevated pKa values meaning there is significant charge regulation upon fibril formation^19^. The aggregation process follows a nucleation-dependent mechanism, which starts with the assembly of two or more monomers^35^. These species, referred to as oligomers, are very transient, and have a higher tendency to dissociate back into monomers than to convert and nucleate fibrillar structure^36,37^. After nucleation, the species formed are more prone to grow than to dissociate, and subsequent secondary nucleation leads to the autocatalytic nature of the aggregation process. Interestingly, oligomers have been shown to be the source of toxicity of αSyn^38,39^, as has been the case for other amyloid proteins^40–43^. However, their heterogeneity, transient nature, and low concentration relative to monomers and fibrils make oligomers challenging to study^37,44–46^.

One of the many methods commonly used for oligomer studies is cross-linking^47^. If the cross-linking reaction is rapid enough, one can capture the transient species by forming covalent bonds in non-covalent complexes. Photo-induced cross-linking of unmodified proteins (PICUP)^48^ has been used to study amyloid oligomers of insulin^49^, tau^50^ or amyloid β^51^, among others. PICUP is a redox reaction that is initiated by exposing the metal complex ruthenium (II) tris-bipyridilcation (Ru(bpy)) to light. The excited state of Ru(bpy) donates an electron to ammonium persulfate (APS), which turns into its oxidant form. This attracts an electron from a nearby amino acid to form a reactive radical, which subsequently reacts with another residue to form a single covalent bond^48^. Being a photo-induced cross-linking method, PICUP is easier to control compared to other chemical cross-linking methods, and can be achieved in considerably shorter times, having been used efficiently with reaction times even below one second^48,51–55^. As a redox reaction, it has the potential to react with different amino acids, but it has shown to have a strong preference for tyrosine and tryptophan side chains^49,51,53,56^. αSyn contains four tyrosine residues at positions 39 (Y39), 125 (Y125), 133 (Y133) and 136 (Y136) (Figure 1), making it a suitable protein to study with PICUP^55,56^. Finally, the fact that it forms a single covalent bond means that the amino acids must be within covalent bonding distance during the lifetime of the radical to be cross-linked. This makes PICUP a highly valuable tool for gaining information on the structure of cross-linked species.

Recently, our group has built a reaction chamber to optimize the method and observe cross-linking at reaction times as short as 1 ms^55^. Using different experimental conditions, we demonstrated the PICUP product of αSyn in solution to be the consequence of oligomer formation, and not of diffusion into close proximity. In the current study, we use this knowledge to study the transient interactions between αSyn monomers in different states. When αSyn is free in solution, it interacts with its neighboring proteins in three dimensions (Figure 1A). When bound to membranes, monomers are sitting on a two-dimensional plane (Figure 1B). Finally, when aggregated into fibrils, αSyn monomers are stacked in a one-dimensional structure (Figure 1C). By performing PICUP of αSyn in these different states, we aim to elucidate the effect the dimensionality of the system on the cross-linking of the protein. Using a combination of various tyrosine to phenylalanine mutants to turn off the cross-linking capacity of different residues in αSyn, we identified the main participants of PICUP cross-linking in the different systems. This has allowed us to identify two main groups of cross-links in solution: Y39-Ct, which happens internally, representative of the N-C termini interaction in monomers in solution; and inter-monomer Ct-Ct, which is the main source of PICUP-visible oligomers. When adsorbed to membranes, the internal cross-linking is blocked while the Ct-Ct cross-linking between monomers remains, presumably due to the high packing density of C-termini sticking into solution. Finally, in fibrils, αSyn can only be cross-linked via Ct to neighboring monomers, reducing its cross-linking severely. This study sheds light into the type of species PICUP can capture, and shows its potential as a method for studying transient interactions of αSyn in different systems.

## 2. Methods

### 2.1. α-Synuclein Expression, Purification and Preparation

Wild-type (WT) α-Synuclein (αSyn), as well as its mutant variants, was expressed in *E. coli* using a Pet3a plasmid with *E. coli*-optimized codons (purchased from Genscript, Piscataway, New Jersey). The protein was purified using heat treatment, ion-exchange, and gel filtration chromatography, as previously described by *Ortigosa-Pascual et al*^19,55^. Each purified protein was aliquoted, freeze-dried and stored at −20° C.

All experiments started with dissolving the aliquot of the protein variant of choice in 6 M GuHCl, followed by a gel filtration on a 10×300 mm Superdex75 column (GE Healthcare). This step serves to ensure fully monomeric sample, and to exchange the buffer to 10 mM MES, pH 5.5, the initial conditions used for all our experiments.

The concentration of WT αSyn was determined by the absorbance at 280 nm and using an extinction coefficient of ɛ = 5800 M^-1^ cm^-1^. Given that the different Tyrosine content of the mutants affects their 280 nm absorbance, but the number of primary amines in the protein remains the same, the concentration of all mutants was determined using *o*-phthalaldehyde (OPA) fluorescence intensity, using WT αSyn for the standard curve^57,58^.

### 2.2. SUV preparation

Lyophilized lipids 1,2-dioleoyl-sn-glycero-3-phospho-L-serine sodium salt (DOPS) and 1,2-dioleoyl-sn-glycero-3-phosphocholine (DOPC) were purchased from Avanti Polar Lipids (Alabaster AL) and stored at -20° C.

Small unilamellar vesicles (SUVs) were formed by extrusion as follows. The required amount of DOPC and DOPS was weighted and mixed in a glass vial in a molar ratio of 7 to 3 and then dissolved in chloroform:methanol (9:1). The organic solvent was then evaporated by applying a gentle flow of nitrogen, and the vial was kept in a vacuum chamber overnight to remove any excess solvent from the lipid film. Lipids were hydrated in 10 mM MES at pH 5.5 and vortexed for a few minutes to get a final lipid concentration of 8 mM. The lipid dispersion was extruded 23 times using an Avanti Mini Extruder (Avanti Polar Lipids) and 100 nm pore size filters. The filters were previously saturated with the same lipid mixture, which was discarded prior to the extrusion process. The size distribution and polydispersity index of the obtained SUVs were analyzed through Dynamic Light Scattering (DLS) using a Malvern Zetasizer Nano-Z (Malvern Instruments). For all the samples used, the average hydro-dynamic diameter and polydispersity index were in the range 92-107 nm and 0.04-0.09 respectively.

### 2.3. Circular Dichroism

Circular dichroism (CD) spectra were recorded using a JASCO J-715 CD spectrometer equipped with a Peltier (PT-348WI) type cuvette holder at 20°C, using 1 mm path length quartz cuvette (Hellma Analytics 110-QS). The CD spectra were recorded using a bandwidth of 5 nm, a data pitch of 1 nm, a continuous scanning mode with speed of 20 nm/min and a response time of 2 s. For each sample, CD spectra were recorded between 250-190 nm three times and averaged. The CD signal originating from the buffer (10 mM MES at pH 5.5) was then subtracted to the CD signal of the protein. The protein concentration in all samples was kept constant at 5 μM while the lipid concentration was varied accordingly for reaching the desired lipid-to-protein ratios (L/P). The CD signal was converted to mean residue ellipticity (MRE) and analyzed as function of the L/P.

### 2.4. Aggregation kinetics

Aggregation of αSyn was monitored with thioflavin T (ThT) fluorescence. 20 µM αSyn (in 10 mM MES, pH = 5.5) were mixed with 3 µM ThT, and aliquoted into a 96-well half area non-treated polystyrene plate (3880 Corning®). The plate was sealed to avoid evaporation and incubated at 37° C without shaking in a FLUOstar Omega plate reader (BMG Labtech, Offenburg, Germany). The aggregation was monitored by measuring excitation and emission wavelengths of 448 and 480 nm respectively, and with the cycle-time adjusted to the minimum-cycle-time to ensure constant mild agitation^59^. Aggregation was followed until the fluorescence reached a plateau.

### 2.5. PICUP

Photo-induced cross-linking of unmodified proteins (PICUP) was performed as described^55^. The sample consisted of 20 µM αSyn in 10 mM MES, pH = 5.5, either by itself, or with added SUVs to a final ratio indicated for each experiment. 18 µL of that sample were then mixed with 1 µL Ru(bpy) (1 mM) and 1 µL ammonium persulfate (APS) (20 mM). The mixture was exposed to 450 nm light for the desired time using a custom-made reaction chamber and stopped by addition of a 5x concentrated gel-loading dye^55^. Samples were analysed by sodium dodecyl sulphate polyacrylamide gel electrophoresis (SDS-PAGE) using Novex^TM^ 10-20% Tricine pre-casted gels. PageRuler™ prestained protein ladder was used as a reference. Gels were stained overnight with InstantBlue^TM^ and scanned with an Epson Expression 10000XL scanner.

## 3. Results

### 3.1. Cross-linking in solution

#### 3.1.1. WT αSyn in solution

Photo-Induced Cross-linking of Unmodified Proteins (PICUP) of α-synuclein (αSyn) in solution leads to the formation of species larger than monomers^55^. This can even be observed in samples where monomeric αSyn has been isolated, due to the fast formation of PICUP-reactive oligomers^55^. The outcome of cross-linking depends on the reaction time, which is controlled by the light exposure to the sample (Figure 2). This way, when the sample is exposed to light for a short duration (i.e., 10 ms), only one new band appears in the sample, corresponding to the size of a dimer (Figure 2A). Increasing the reaction time increases the chances of cross-linking bigger species. However, it also has the risk to cross-link species more than once, either between monomers or internally within the monomer, leading to products with more complex morphologies. This causes the formation of more than one band per oligomer size. When many bands with slight variation in electrophoretic mobility are formed, the individual bands merge together to form a diffuse long band, as seen when doing the reaction for 1s or more (Figure 2A).

**Figure 2.**
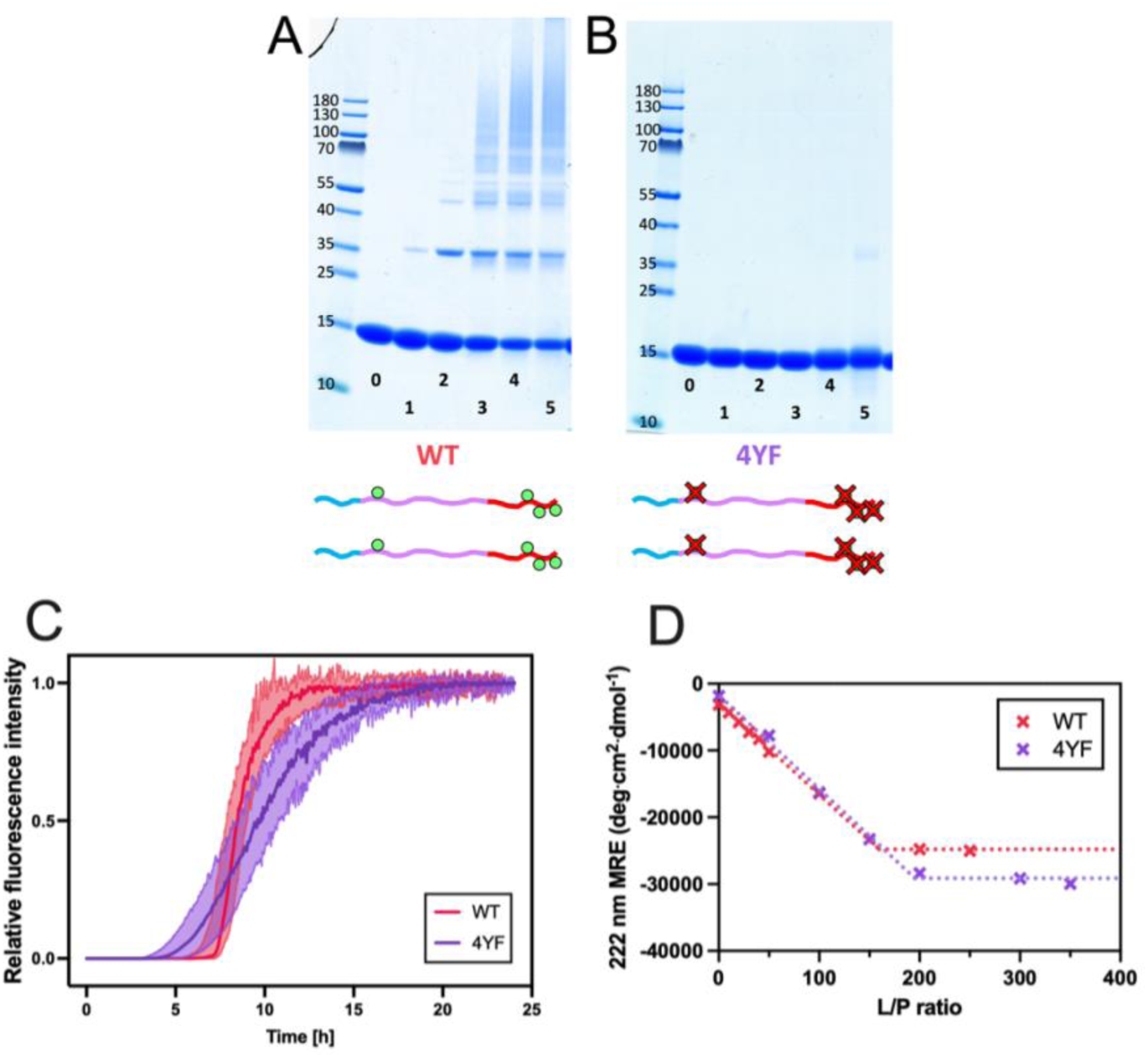
Role of tyrosine residues in aSyn PICUP, aggregation and lipid binding. PICUP of WT (A) and 4YF (B) αSyn in solution. The reaction was performed for 0 ms (0), 10 ms (1), 100 ms (2), 1 s (3), 10 s (4) and 100 s (5). Full gels can be seen in Figure S1. (C) Aggregation kinetics of WT (red) and 4YF (purple) αSyn monitored by ThT fluorescence. (D) Lipid binding of WT (red) and 4YF (purple) αSyn measured by CD. Mean residue ellipticity (MRE) at 222 nm was measured as a function of L/P ratio for both proteins (cross marks). The dotted line shows the trend of the MRE as a function of L/P, with the intersection indicating the L/P value at which no more α-helix is formed (WT ≍ 150, 4YF ≍ 200).

#### 3.1.2. Tyr → Phe mutation

Previous studies have identified tyrosine (Tyr, Y) as the main source of cross-linking of αSyn via PICUP^51–53,56^. Furthermore, phenylalanine (Phe, F), retaining the aromaticity but lacking the hydroxyl group of Tyr, seems to have a way lower efficiency of cross-linking^52,60^. With the aim to eliminate the cross-linking capacity of αSyn, we therefore expressed a mutant form of the protein with the four Tyr residues (Y39, Y125, Y133 and Y136) mutated to Phe, named 4YF.

When doing PICUP with the 4YF mutant under the same conditions as for the wild type (WT) αSyn, we indeed observed close to zero cross-linking (Figure 2B). A very faint band can be seen when the reaction is performed for as much as 100 s, which can thus be attributed to the very low efficiency of non-tyrosine cross-linking. In order to evaluate whether the lack of cross-linking of 4YF is due to a drastic change in protein behaviour, the mutant protein was subjected to aggregation and vesicle adsorption studies (Figure 2C & 2D). The ThT fluorescence intensity as a function of time shows that 4YF aggregates on a similar timescale to WT αSyn (Figure 2C), albeit the more shallow transition of the mutant may reflect a smaller dominance of secondary nucleation^61^. When it comes to adsorption of the protein monomer to SUVs, up to the first 100 residues of αSyn undergo a conformational change from a disordered to an α-helical structure^27,62,63^. This transition can be probed by means of circular dichroism (CD) spectroscopy, which provides an estimation of the L/P ratio at which all proteins are bound to the SUVs, here referred to as saturation point. Figure 2D shows that the adsorption behaviour of the 4YF mutant to the SUVs is similar to that of the WT αSyn, with saturation points of L/P≍150 and L/P≍200 for the WT and 4YF proteins, respectively. Taken together, these results indicate that the observed differences in ability of the 4YF mutant to cross-link are due to the exchange of tyrosine to phenylalanine rather than changes in the overall self- and co-assembly properties of the protein caused by the mutations.

#### 3.1.3. Role of each tyrosine in αSyn PICUP

To determine the role of each Tyr residue in the cross-linking of αSyn in solution, four different single amino-acid Tyr → Phe mutants were purified and used in PICUP studies in the same manner as the wild-type: Y39F, Y125F, Y133F and Y136F. The three C-terminal mutants (125, 133 and 136) showed the same behavior as WT αSyn, with the formation of a diffuse long band under longer reaction times (Figure 3B, 3C & 3D).

**Figure 3.**
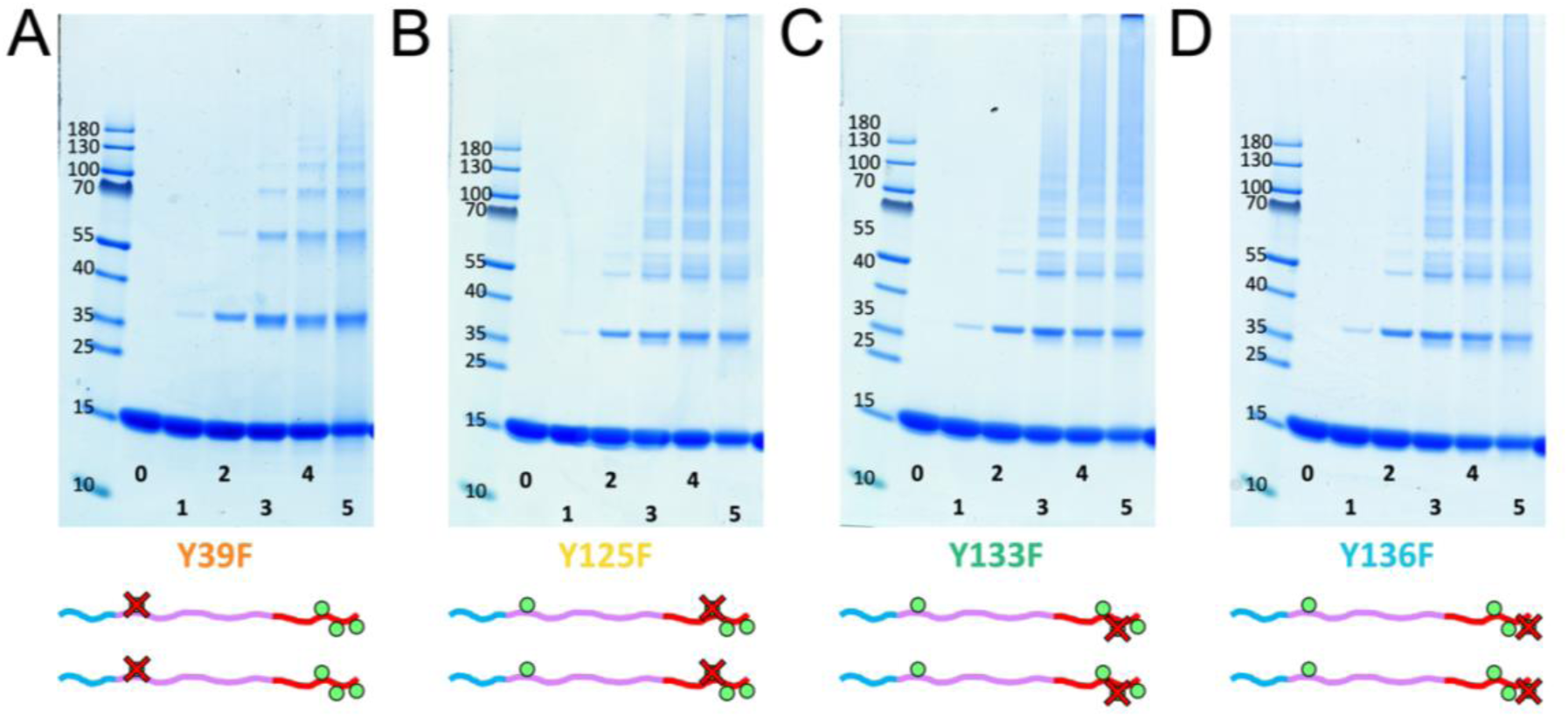
PICUP of αSyn with single Tyr→Phe mutations. PICUP was performed for αSyn mutants Y39F (A), Y125F (B), Y133F (C) and Y136F (D) to evaluate the effect of Tyr → Phe mutations in PICUP. The reaction was done for 0 ms (0), 10 ms (1), 100 ms (2), 1 s (3), 10 s (4) and 100 s (5). Full gels can be seen in Figure S2.

However, Y39F shows a strikingly different pattern (Figure 3A). Increasing the reaction time of Y39F leads to the formation of new bands of higher MW, but not to the point where the bands blend into a diffuse long band. Rather, the results reveal a single band for each oligomer size. This points at the presence of Tyr39 being responsible for a higher variation of cross-linked products in the WT αSyn.

To further evaluate the role of Y39, a fifth mutant was expressed, with Y125, Y133 and Y136 all mutated to F. This mutant, named 3YF, contains Y39 as the only potential source of cross-linking. The cross-linking of this mutant led to the formation of dimer bands only, and only after a very long reaction time (100 s) (Figure 4A).

**Figure 4.**
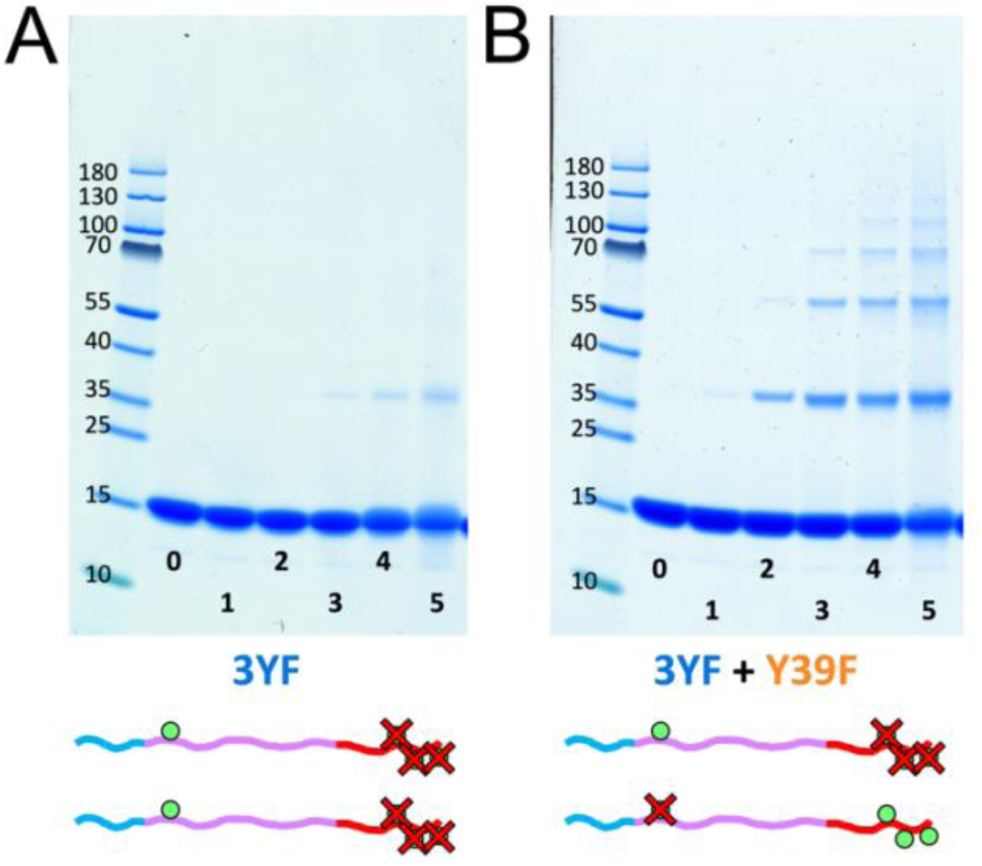
Internal vs monomer-monomer cross-linking. (A) PICUP of 3YF mutant of αSyn, having the three C-terminal Tyr (Y125, Y133 and Y136) mutated to Phe. (B) PICUP of a 50:50 mixture of αSyn mutants 3YF and Y39F. The reaction was done for 0 ms (0), 10 ms (1), 100 ms (2), 1 s (3), 10 s (4) and 100 s (5). Full gels can be seen in Figure S3.

#### 3.1.4. How does Y39 contribute to heterogeneity of PICUP products?

Y39 could be leading to a higher variation of cross-linked morphologies either by participating in monomer-monomer cross-links, or by leading to internal cross-linking within a monomer. To distinguish between these two possible contributions, we prepared a reaction with a 50:50 mixture of Y39F and 3YF. As these mutants contain either the C-terminal tyrosines or Y39, respectively, the ability of the monomer to internally cross-link is blocked, while allowing for monomer-monomer cross-linking between the Y39 and the C-terminal tyrosines. Strikingly, cross-linking of this sample led to the formation of bands akin to those observed for Y39F alone (Figure 4B).

### 3.2. Cross-linking of protein adsorbed at lipid vesicles

While proteins in bulk interact with each other in 3D, the adsorption of αSyn to a lipid mebrane surface places the proteins in a 2D plane, which alters their potential interactions. When αSyn is cross-linked in the presence of SUVs, the outcome changes drastically depending on the relative concentration of protein and SUVs in terms of the lipid to protein ratio (L/P) (Figure 5A). Increasing the L/P ratio leads to the formation of more defined bands. This may be due to both a transition to a 2D system, or a change in structure of the protein when adsorbing to the lipid vesicle. The change to more defined bands from solution to lipid bound becomes most pronounced at L/P ≥ 150, which happens to be the saturation point for WT αSyn (Figure 2D). This strongly indicates that the bands observed at both L/P = 150 and 250 represent the cross-linking of fully adsorbed αSyn, and that the transition we observe between L/P = 0 and L/P = 150 can be related to the proportion of proteins that is free in solution and the ones adsorbed to the SUVs.

**Figure 5.**
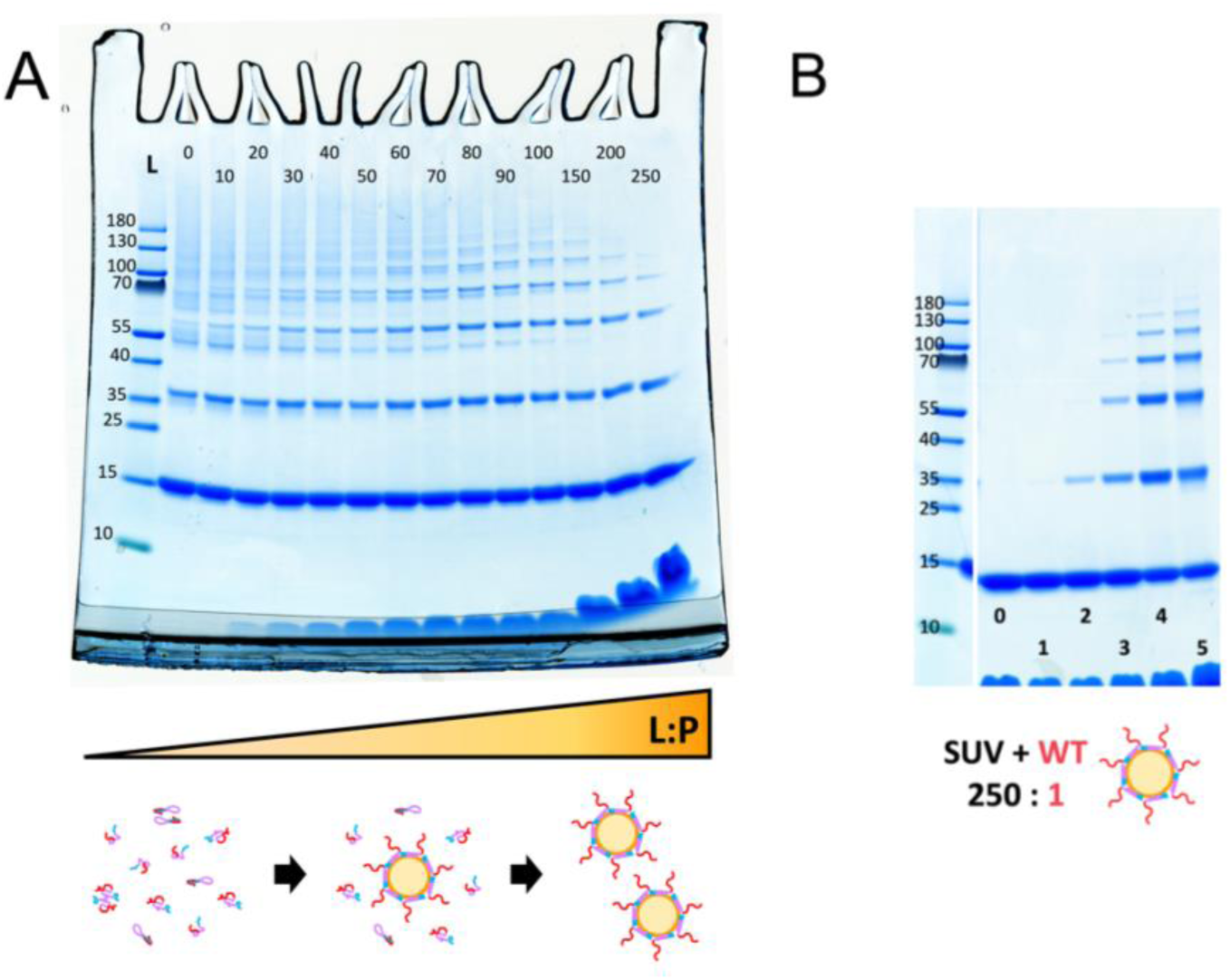
Effect of lipid binding on PICUP of αSyn. (A) PICUP was performed to αSyn for 1 s in the presence of SUVs at different L/P ratios (0 to 250). (B) Using a L/P = 250 where we know all the αSyn will be bound to the SUVs, PICUP was performed for 0 ms (0), 10 ms (1), 100 ms (2), 1 s (3), 10 s (4) and 100 s (5). Full gel of figure B can be seen in Figure S1.

To study the protein when fully bound to the SUVs, we used a system with a L:P ratio of 250, which is above the saturation points of both WT and 4YF (Figure 2D), meaning that no further binding occurs even if more vesicles are added. Performing PICUP at different lighting times for WT at L/P = 250 shows that increasing the lighting time doesn’t generate new bands other than the well-defined ones (Figure 5B). Strikingly, these well-defined bands are of the same molecular weight as those observed for the Y39F mutant in solution (Figure 3A). This is more clearly seen in the full gels, available in supplementary information (Figures S1 and S2).

The same experiments at saturating L/P (L/P = 250) were run for the different mutants (Figure 6). To evaluate the size and distribution of bands obtained for each mutant in the presence of SUVs, WT αSyn was cross-linked for 10 s in presence of SUVs at a L/P = 250, and used as a reference. All the single point mutants, regardless of their cross-linking in solution, displayed the same behavior as that of WT αSyn when adsorbed to vesicles. Interestingly, vesicle-adsorbed Y125F seems to show less bands than WT αSyn under the same conditions (Figure 6B). Mutants 4YF and 3YF were also evaluated under the same conditions (Figures S1 and S3). 4YF showed absolutely no bands when adsorbed to SUVs, and 3YF behaved the same as in solution, forming only dimers.

**Figure 6.**
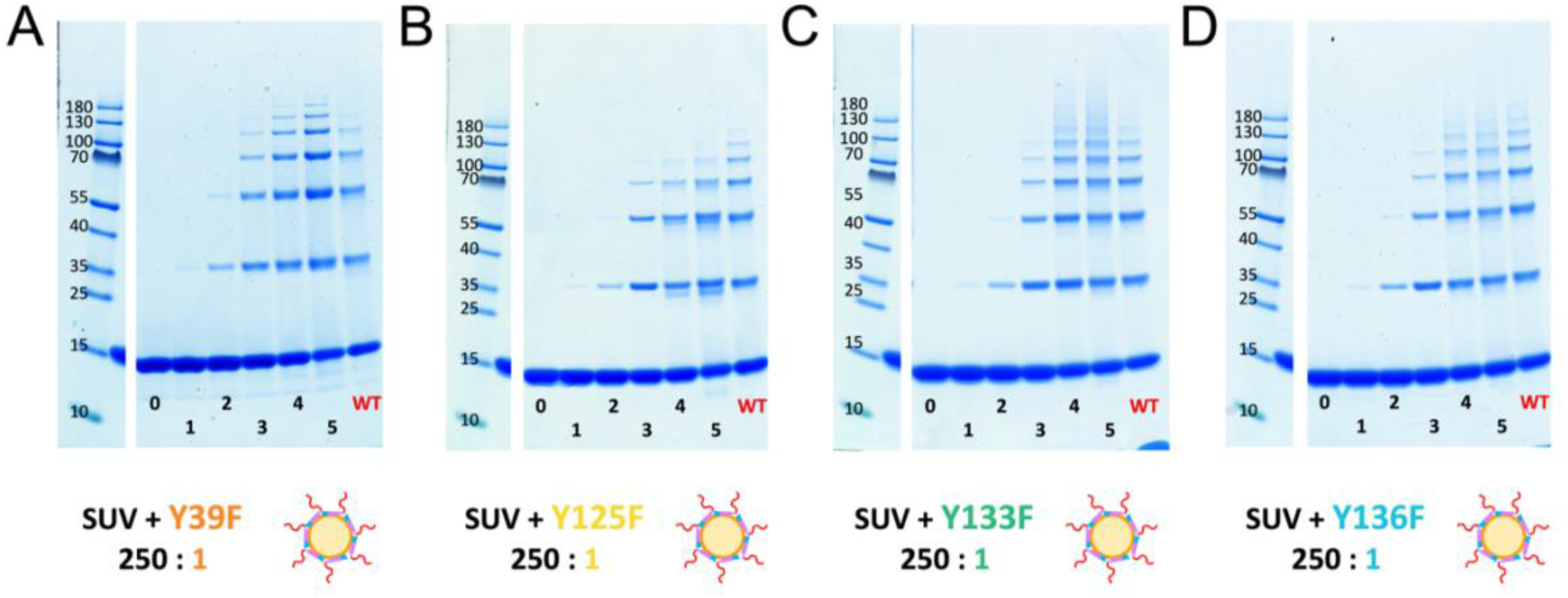
PICUP of single Tyr→Phe αSyn mutatants. PICUP was performed for αSyn mutants Y39F (A), Y125F (B), Y133F (C) and Y136F (D) in the presence of SUVs at a L/P ratio of 250. The reaction was done for 0 ms (0), 10 ms (1), 100 ms (2), 1 s (3), 10 s (4) and 100 s (5). Fnally, a WT αSyn at L/P = 250 was cross-linked for 10 s and loaded in all gels as a reference (labelled WT, in red). Full gels can be seen in Figure S2.

### 3.3. Cross-linking in fibrils

When αSyn aggregates into fibrils, the monomers stack together forming β-sheets parallel to the fibril axis. This way, the αSyn monomers are essentially stacked in 1D, reducing the available interaction partners of each monomer from the in solution and lipid adsorbed systems. As previously reported, performing PICUP on fibrillated αSyn leads to the formation of almost no bands^55^. In fact, when PICUP is performed at different times during the aggregation process, the cross-linked species can be seen decreasing in concentration (Figure 7A). This observation applies to all the Tyr → Phe mutants too (Figure 7B). All single point mutants (Y39F, Y125F, Y133F & Y136F) show a behavior that is undistinguishable from that of WT αSyn. As previously reported, two bands below the cross-linked dimer band can be observed upon aggregation of WT αSyn, which are independent of cross-linking^55^. This same phenomenon is observed for all mutant αSyn variants (Figure 7B).

**Figure 7.**
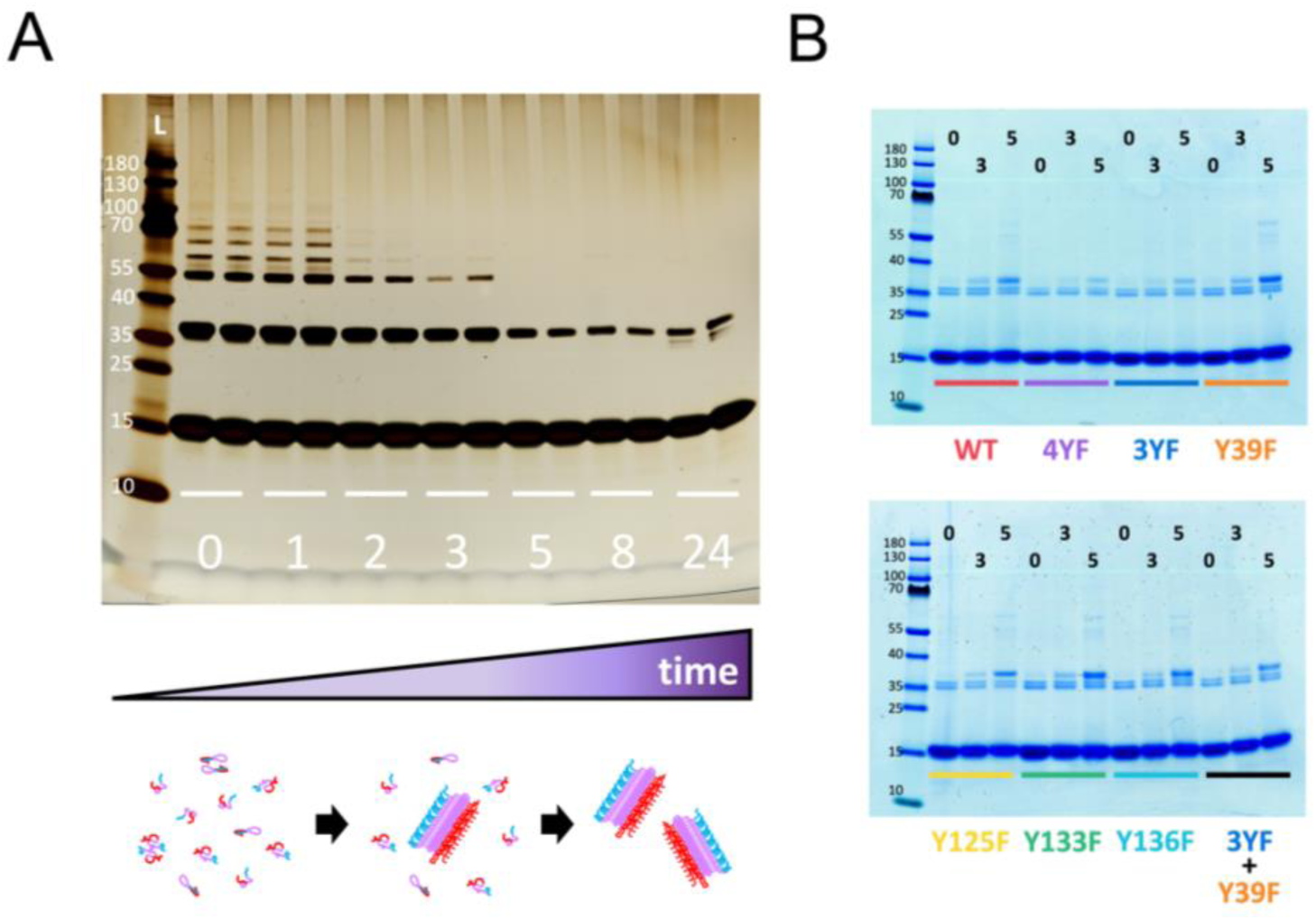
PICUP of αSyn on fibrils. (A) PICUP at 1s lighting time of WT αSyn over aggregation time. The numbers indicate the time of aggregation in hours. Sample was collected before the start of aggregation (0), middle of lag phase (1), start of exponential phase (2), middle of exponential phase (3), end of exponential phase (5), final plateau (8), and late after full aggregation (24). Adapted from Ortigosa-Pascual et al. Figure 7, panel D ^55^ (B) PICUP of fully fibrillated WT αSyn as well as all the Tyr→Phe mutants of αSyn used in this study. Sample was cross-linked for 0 ms (0), 1 s (3) and 100 s (5).

When all Tyr are removed (mutant 4YF), the cross-linked dimer band becomes weaker, but it does not completely disappear, suggesting a non-Tyr amino acid is cross-linking in fibril form too. In fact, when comparing 4YF and WT αSyn, the reduction of cross-linking product caused by the removal of all Tyr is much lower in fibrils than it is in solution. The fact that 4YF and 3YF show a similar decrease in cross-linked dimer band moreover points at the possibility of Y39 not taking part in cross-linking in fibrils.

## 4. Discussion

Studies of the self- and co-assembly of αSyn are crucial in order to fully understand its native and pathological functions. Here, we show the potential and versatility of PICUP as a tool to study transient interactions between αSyn molecules in different environments. We find that the dimensionality of the system studied alters the cross-linking pattern, and parallel studies in different geometries thus provide more information on the relative positioning of αSyn (Figure 9).

### 4.1. Cross-linking in solution

As observed in an earlier study^55^, PICUP of αSyn in solution depends highly on the lighting time used to trigger the reaction. The presence of clear bands of defined size, in combination with diffuse long bands, suggests the coexistence of at least two groups of cross-linked species and reflects on the complexity of the cross-linking that occurs upon long reaction-time. By mutating Tyr to Phe, we aimed to turn off individual tyrosines to help us understand the cross-linking pattern.

#### 4.1.1. Tyr → Phe mutations decrease αSyn PICUP reactivity

Substituting all Tyr residues with Phe (mutant 4YF), while keeping aggregation and lipid adsorption properties similar, severely decreased the cross-linking of αSyn. The presence of a faint band at 100 s lighting time suggests that a residue other than tyrosine has the ability to cross-link, but with very low reactivity. This could be a residue present in WT αSyn, or the newly added Phe residues, which do show weak cross-linking reactivity^52^.

When the three Tyr residues of the C-terminus were replaced (3YF), leaving only Y39 available to cross-link, only dimer bands were observed. These bands show higher intensity than those observed for the 4YF mutant, suggesting Y39 being involved in the formation of these cross-linked species. The fact that a variant with a single Tyr residue can only form dimers implies that a Tyr cannot cross-link more than one other Tyr at a time. From a chemical perspective, a dityrosine could still react with a third Tyr and form a trityrosine^64^. However, given the tight distance restriction of PICUP, the two Tyr that become cross-linked must be within a covalent bond distance during the lifetime of the radical. This is of course more hindered for a more bulky dityrosine, which also has its mobility severely restricted by the first covalent bond, making a triple-Tyr cross-link highly unlikely.

#### 4.1.2. Y39 is responsible for diffuse bands in PICUP of αSyn

When the four Tyr residues of αSyn were mutated to Phe one at a time, a clear difference could be observed between Y39 and the three Tyr in the Ct. While mutating either of the Ct residues did not lead to any detectable changes in the outcome of PICUP, the mutation of Y39 led to the disappearance of the longer diffuse bands. This indicates that Y39 is the responsible for increasing the complexity of the cross-linking pattern, leading to a higher number of bands.

Figure 8 illustrates the different ways the addition of Y39 can increase the complexity of the species resolved by SDS-PAGE. In this figure, each protein is represented as a line. Given the proximity of residues Y125, Y133 and Y136, we expect proteins cross-linked via any of them to be similar enough in morphology to be indistinguishable by SDS-PAGE. Due to that, and for the sake of simplicity, we will refer to the three of them together as cross-linking the Ct. Thus, there are three possible cross-links between monomers: Y39-Y39, Y39-Ct and Ct-Ct.

**Figure 8.**
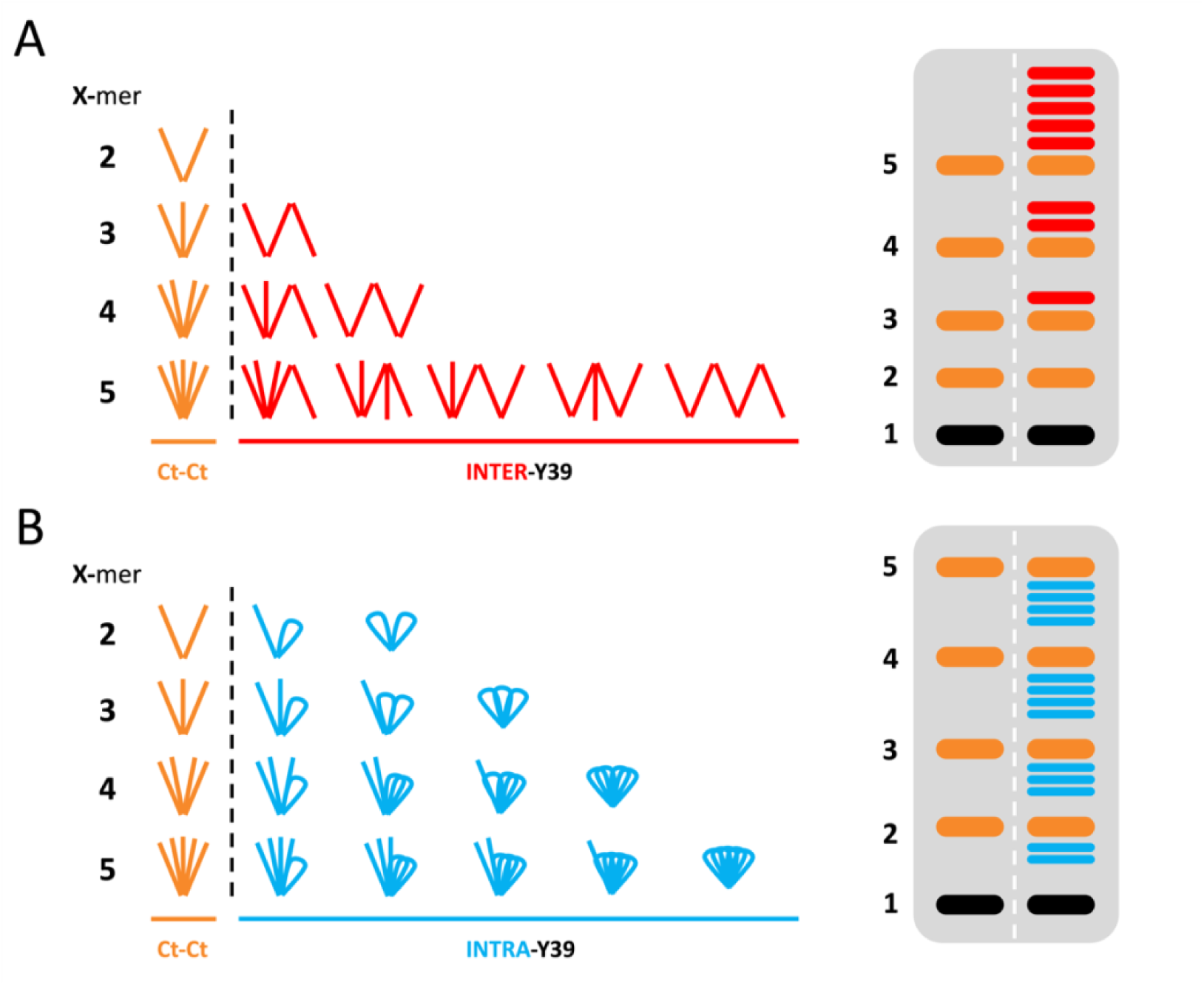
Schematic representation of the effect of Y39 in cross-linking of αSyn. Each line represents an αSyn monomer. The system is simplified in such way that Y125, Y133 and Y136 are all reduced to a single cross-linking site (Ct). On top of that, Y39 and Ct cross-linking are treated as symmetrical cross-links. This way, if the only cross-linking occurring via PICUP is that between C-termini, we would expect each oligomer size to give rise to a single morphology, labelled “bouquet” morphology (“Ct-Ct”, coloured orange). If Y39 is involved in inter-molecular cross-linking (A), new morphologies may be generated (“INTER-Y39”, red). The number of possible morphologies increases with the size of the oligomer. We expect a more branched-out structure to migrate more slowly than the Ct-Ct (bouquet) oligomers of the same order. If the Y39 was instead involved in Intra-molecular cross-linking (B), it would cause the “folding” of a monomer onto itself (“INTRA-Y39”, blue). This would give rise to new morphologies, which increase in number the bigger the oligomer order. As these structures would be more compact than the ones in the bouquet morphology, we would expect them to migrate faster than the Ct-Ct oligomers of the same order. Differentiation between Ct and Nt would lead to an even bigger difference between systems, as is reflected in Figure S4 for a system with inter-Y39 cross-linking.

Based on these simplifications, if Ct-Ct cross-links only occur in the system, we would expect a single morphology, and thus, a single band per oligomer size (i.e., one for dimer, one for trimer, etc.). All the monomers in these oligomers would be bound together via Ct in what we call a “bouquet” morphology (labelled “Ct-Ct”, coloured orange). The addition of Y39 to the system could increase the complexity of αSyn cross-linking in three ways. Firstly, if Y39 participated in inter-molecular cross-linking (Figure 8A), the combination of Ct-Ct, Y39-Y39 and Ct-Y39 cross-links would lead to the formation of additional morphologies (labelled “INTER-Y39”, coloured red). These structures, being capable of expanding to longer species than the “bouquet” oligomers, would be expected to migrate less than the Ct-Ct oligomers. Secondly, if Y39 participated in intra-molecular cross-linking (Figure 8B), that would cause the monomers to fold over themselves, creating oligomers with a different migration (labelled “INTRA-Y39”, coloured blue). These oligomers, being more compact than the “bouquet” oligomers, would be expected to migrate further than the Ct-Ct oligomers. Thirdly, Y39 could be contributing to the cross-linking pattern by participating in both inter- and intra-molecular cross-linking.

This way, the system with only Ct-Ct cross-linking would lead to a 1 oligomer = 1 band pattern. However, if Y39 is involved in either inter- or intra-molecular cross-linking, it would lead to 1 oligomer ≥ 1 band. In either case, the bigger the oligomer, the more possible morphologies of the cross-linked species, explaining why the diffuse behaviour observed at high lighting times is more prominent at higher molecular weights. The morphologies represented in Figure 8 would be subdivided into more if we distinguish between Nt and Ct. This would increase the complexity of the system even further, as cross-linking between N-termini or between C-termini could affect the morphology (Figure S4).

#### 4.1.3. Y39 participates in intra-molecular Y39-Ct binding only

By mixing mutants Y39F and 3YF, we create a system where Y39-mediated inter-molecular cross-linking is allowed, thus having the possibility of forming Y39-Y39, Ct-Ct, and interY39-Ct cross-links. However, intraY39-Ct cross-linking cannot occur. Strikingly, PICUP of the Y39F and 3YF mixture looks exactly the same as that performed with Y39F alone. This means that removing the ability of Y39 to do intra-molecular cross-linking has the same effect as removing the ability of Y39 to cross-link at all. This finding proves that the only role of Y39 in the cross-linking of WT αSyn is intra-molecular Y39-Ct binding (intraY39-Ct). Additionally, this means that during PICUP of WT αSyn no inter-molecular Y39-Ct nor Y39-Y39 cross-linking form, as those would have led to the formation of more bands than observed for Y39 alone. In short, the only contribution of Y39 to the cross-linking of αSyn is the one depicted in Figure 8B.

Given the observation of cross-linked dimers with the 3YF mutant, attributed to Y39-Y39 cross-linking above, it would seem that Y39-Y39 can form in absence of other cross-linkable residues, but not when the three Ct Tyr are present. The fact that 3YF needs more light to form dimers than WT and mutants containing the Ct Tyr do, suggests Y39-Y39 to be a less favoured cross-link via PICUP. In addition, the Y39-Y39 cross-link might not appear to form in WT αSyn for several reasons. Firstly, intraY39-Ct may have a strong preference over Y39-Y39. In addition, Y39-Y39 cross-links in WT αSyn may occur only between monomers that already contain a Ct-Ct cross-link, which would not alter the migration of the oligomer.

The obtained results imply that two cross-links are preferentially formed in solution: intraY39-Ct and Ct-Ct (Figure 9). These cross-links are most likely favoured by transient interactions within and between αSyn monomers.

**Figure 9.**
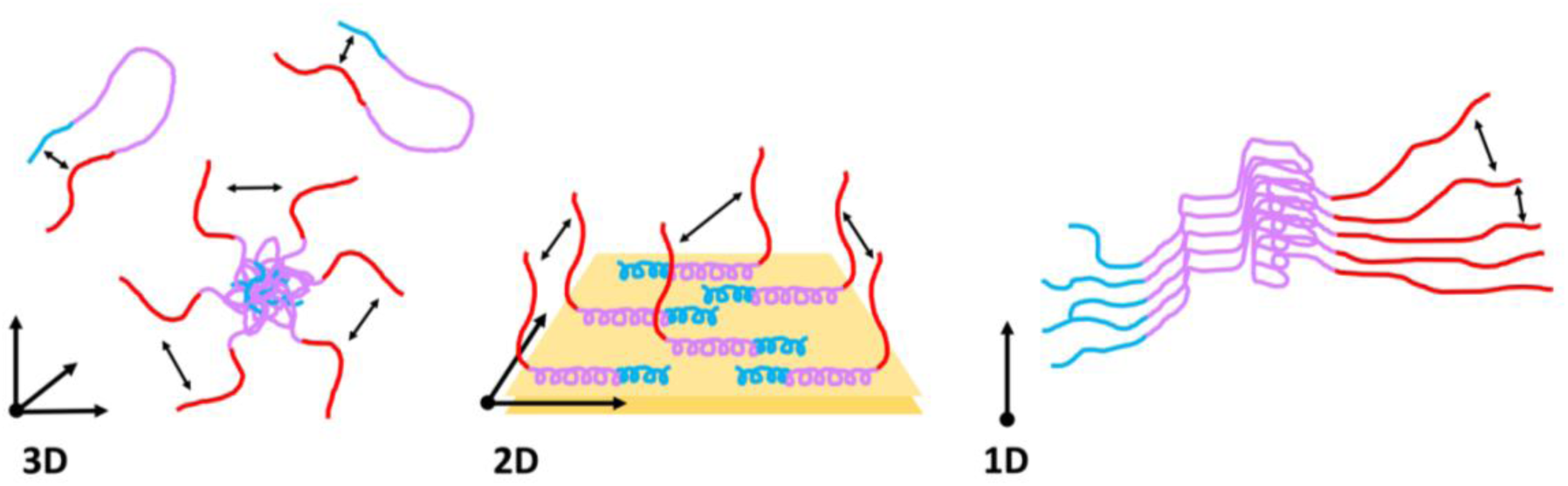
Cartoon of transient interactions of αSyn detected with PICUP, and their dependence on the dimensionality of the system. In solution (3D), PICUP reports on internal Ct-Nt interactions, as well as Ct interactions of oligomeric species. When adsorbed to lipids (2D), only Ct-Ct interactions are detected between proteins. When αSyn is fibrillated, only the Ct-Ct cross-linking is detected, albeit with less frequency than in the other two systems.

#### 4.1.4. IntraY39-Ct cross-linking: αSyn’s self-chaperoning Ct

The presence of intraY39-Ct can only come from an interaction between the N and C termini of αSyn. Previous studies have shown that monomeric αSyn adopts a more compact conformation than that of a random coil, due to long-range interactions between its Nt and Ct^65,66^. Internal Nt-Ct attraction may disfavour the interactions between NAC regions of different monomers, preventing the protein from aggregation^21,23,65–69^. Binding of Ca^2+^, Cu^2+^ and polyamines to the Ct may expose the Nt and NAC domain, speeding up aggregation^70–72^. Truncations of Ct, which occur *in vivo* both in healthy and diseased brains^73–76^, lead to αSyn adopting a more extended conformation^66^. Truncation of Ct leads to a faster fibril formation^23,77–82^, higher toxicity^83–85^ and increased mitochondrial damage^66^. On top of that, Ct truncation increases the interaction of αSyn with lipid membranes and molecular chaperones by freeing the Nt involved in those interactions^66,86^. In short, the Ct plays a self-chaperoning role in the self- and co-aggregation of αSyn by interacting with the Nt.

The finding that intraY39-Ct is one of the two main cross-links obtained with PICUP of αSyn would be expected based on previous finding of interactions between Nt of monomers and the exposed Ct of fibrils. The screening of the monomer-fibril interaction by salt as seen in experiment and simulations imply that electrostatic interactions are the main driving forces^21^. However, it has been discussed that not only electrostatics, but π-π interactions between aromatic residues in both ends also play a key role in the Nt-Ct interaction^87^. Internal Nt-Ct interactions as observed by NMR and in simulations^21,22^, imply that Y39 and Ct will be in close enough proximity for PICUP to cross-link them. Furthermore, dityrosine Nt-Ct cross-linked monomers obtained via oxidation have been shown to inhibit αSyn aggregation via steric hindrance^88^. PICUP cross-linked oligomers have also shown the same effect on aggregation^56^, which can now be explained with intraY39-Ct cross-linking. The finding of internal Y39-Ct cross-linking reflects on the key role of Ct in monomeric αSyn and shows PICUP to be a very useful tool to monitor this phenomenon.

#### 4.1.5. Ct-Ct cross-linking: oligomeric αSyn’s interactions

##### InterY39-Ct occurs first, but Ct-Ct is representative of PICUP-oligomers

The Ct-Ct cross-linking must reflect the formation of aggregated species, e.g. oligomers. Given the proximity between the Nt and Ct in transent conformations of a monomer, the Y39-Ct interaction may occur earlier than the Ct-Ct one. However, mutants without the ability to form internal Nt-Ct cross-linking also show oligomeric bands (Figure 3A, Figure 4B), suggesting that Ct-Ct crosslinks do not depend on intraY39-Ct cross-linking. Thus, the oligomers captured via Ct-Ct are not artifacts due to a prior internal cross-link of monomers. This means that the proximity between C termini is a characteristic feature of PICUP-visible oligomers (PvO).

##### Previous reports on αSyn oligomer structures

Although heterogeneous, αSyn oligomers are on average more stable than oligomers formed by other amyloids, which can be ascribed to the fact that αSyn oligomers are more structured than other amyloid oligomers^37^. In fact, many studies show them to be rich in β sheets^37,89,90^, most often anti-parallel β-sheets^91–93^. Interestingly, the anti-parallel β-sheet structure is reported for kinetically trapped oligomers (KTO), species generated through very precise protocols and showing a very high stability^92,93^. It has been hypothesised that the anti-parallel β-sheet found in KTO is the reason behind their stability, as it must overcome a high energy barrier to transition into a parallel β-sheet compatible with fibrillar structures^90,93^. Inversely, parallel β-sheet oligomers, being more compatible with the fibrillar structure, are more likely to quickly turn into fibrils and grow, showing a more transient behaviour^93^. Some studies of oligomers have shown Ct truncation of αSyn to cause a decrease in oligomer formation^23^ while others show an increase^66,85^. Regardless of this, Ct is shown to play a key role in oligomer formation. The oligomer structure determined for KTO show a structured core, with an unstructured Ct tail^92,94^. Additionally, while some KTO considered to be *off pathway*^95^ have the Nt immobilized in the oligomer core, the ones showing heavier toxicity have the Nt accessible, this feature being an important part of membrane binding, disrupting and cell toxicity^94^.

##### What structure do our PICUP oligomers most likely have?

In a previous study, we have shown that αSyn PvO appear quickly after monomer isolation, and their concentration dependence follows that of the monomer^55^. These results indicate that αSyn PvO are species in fast equilibrium with monomers. They show low kinetic stability, or persistence, as their concentration is determined by monomer depletion^37^. We here find that the Ct of monomers must be accessible to each other in PvO. These oligomers can be obtained independent of an intraY39-Ct, as seen for WT, but not possible for Y39F. The fact that the intraY39-Ct does not restrict oligomer formation could point towards an oligomer structure that has both N and C termini exposed. Given that Ct-Ct is the main cross-link observed for PvO, it is likelier that the monomers are in a parallel β-sheet, as opposed to the KTO identified in literature, composed mostly of anti-parallel β-sheet^92,93^. Given that the amount of PvO follow the concentration dependence of monomer, and do not seem to be as stable structures as the KTO, it stands to reason that they are species on pathway to fibrillar structure. However, we are only observing a few monomers together. Thus, the observed bands could come either from small oligomers, for which there is no structural information available, or a small fraction of monomers within a bigger oligomer, where all monomers might not have the exact same structure, and therefore our information doesn’t necessarily reflect the structure of the whole oligomer. Performing PICUP for KTO with a more determined secondary structure or combining PICUP with higher throughput methods for structural analysis would help understand the nature of the PvO.

### 4.2. Cross-linking of protein adsorbed at lipid vesicles

When adsorbed to a negatively charged membrane, the first ca. 100 residues of αSyn adopt an amphipathic α-helical structure, which sits with some residues exposed to the solution and others accessing the hydrophobic acyl layer of the membrane^4^. Meanwhile, the Ct tails remain disordered in solution, forming a polymer brush^3^. The results of PICUP for αSyn adsorbed to lipids reflects on this previously described system.

αSyn mutant 4YF shows less bands in lipids than it does in solution, suggesting the residue that cross-links in solution is not in such close proximity in lipids. Helical projections of the α-helix adopted by the Nt of αSyn show Y39 on the upper part of the helix^9,96^, indicating it would be exposed to the solvent, and available for potential cross-linking. 3YF is cross-linked as dimers when adsorbed to SUVs, which indicates that a minimal amount of Y39-Y39 can form in this state. Similarly to what was observed in solution, the formation of a trityrosine is suggested to be structurally restricted also for the membrane-bound protein.

When looking at the effect of adding more membrane (increasing the L/P ratio) on the PICUP of αSyn, we can see that adsorbing to SUVs leads to the reduction of the number of bands, similarly to removal of intraY39-Ct. The bands formed by intraY39-Ct cross-linking in solution (left) disappear as the L/P increases and the internal cross-linking is reduced (Figure 5A). In line with the description in Figure 8B, the bands caused due to intraY39-Ct cross-linking migrate faster than those formed by Ct-Ct alone. For instance, the dimer band in the lipid-free sample has a shadow below it, which becomes weaker upon the addition of membrane and finally disappears at L/P = 150. Similarly, the trimer for the lipid-free sample has two distinguishable bands, one with faster migration at ∼ 55 kDa and one with slower migration at ∼ 60 kDa. As L/P increases, the ∼ 55 kDa band disappears as the other one becomes stronger, until only the ∼ 60 kDa one is visible at L/P = 150. Because the band seen at L/P = 150 reflects saturation, where essentially all protein molecules are bound, and formed via Ct-Ct cross-linking, the band appearing at ∼ 55 kDa must be caused by intraY39-Ct cross-linking of the protein in solution.

This shows that, when adsorbed to lipids, Ct-Ct cross-linking still occurs, while intraY39-Ct cross-linking becomes impossible (Figure 9). Based on the measured L/P at saturation, together with an approximate area per lipid headgroup in the DOPC/DOPS (7/3) bilayer of 70 Å^2^ ^97,98^, we can estimate the average area per protein at the bilayer to ca. 1 protein per 5300 Å^2^. This means the average distance between C-termini would be ca. 73 Å. For a fully extended polypeptide, a length of 7.23 Å per two amino-acids is estimated^99^. Based on that, the remaining 40 amino acids of the exposed Ct would have a length of ca. 145 Å when fully extended, twice the average lateral distance between the points where the C-termini start to extend around residue 100. If we instead calculate the radius of gyration (R_g_) of a random coil, described by Rg = R_0_N^v^, where R_0_ = 1.93 Å and v = 0.60, based on experiments and simulations^100^, and N stands for the number of amino-acid rediues we get R_g_ ∼ 18 Å for a 40-residue segment. This means that there is plenty of room for the Ct tail to fold all the way to come into contact with its own Nt. The lack of observed Y39-Ct crosslinking suggests that charge repulsion may favor a more extended state, in which case the flexible Ct may contact each other by random diffusion, and generate Ct-Ct cross-linking. However, the severe distance restriction of PICUP does not exclude that the cross-linking is reflective of specific interactions between C-termini. Moreover, due to the fluidity of a lipidic bilayer, αSyn proteins are not anchored in space and can move within the membrane, which may facilitate the interactions between the embedded proteins.

The fact that Y125F shows slightly fewer bands than Y133F and Y136F hints towards the possibility of Y125 being the main source of Ct-Ct cross-linking in *α*Syn adsorbed to lipids. Y125 has been previously suggested to be the most PICUP-reactive Tyr residue in the Ct by monitoring of diTyr formation^60^. Further analysis of the role of each amino-acid in the Ct-Ct cross-linking in SUVs has the potential to shed light on the behaviour of lipid-adsorbed αSyn. Finally, since it has been shown that Nt availability is crucial for oligomer activity and toxicity, oligomers with only Ct-Ct cross-linking might be more representative of toxic species^94^. This means that performing PICUP in presence of SUVs could be a useful tool to generate toxic oligomers with WT αSyn, without the intraY39-Ct that occurs in solution.

### 4.3. Cross-linking in fibrils

When performing PICUP for fibrillated WT αSyn, we obtain dimer bands only. 4YF αSyn shows less bands than WT; however, the decrease in cross-linking when comparing WT to 4YF is lower in fibrils than it is in solution. The fact that 3YF shows the same intensity as 4YF suggests that Y39-Y39 cross-linking does not happen for WT αSyn in fibrils. Fibrillar αSyn structures show Y39 next to the fibril core, with the Y39 residues of neighbouring monomers being π-π stacked face-to-face in close proximity to one another^33,101^. This could hinder the rotation of the aromatic rings, limiting the possibility for a covalent-bond formation between neighbouring Y39. Mutants containing at least one of the Ct Tyr residues show as much cross-linking as WT αSyn does, whereas 4YF and 3YF show less. In fibrillar conformation, intraY39-Ct cross-linking cannot occur due to the long distance between the termini^33^. This suggests Ct-Ct cross-linking still occurs between monomers in a fibril (Figure 9). Due to the proteins being stacked in 2D, each monomer has fewer neighbouring monomers to cross-link with, leading to the formation of dimers only, and for some of the mutants a faint trimer band is observed. Based on a previously identified αSyn fibril structure^102^, the residues Lys97 of the monomers in the fibril where the random coil C-termini are anchored, are separated by a distance of ∼ 5 Å. Given the close proximity, we measured very little Ct-Ct cross-linking compared to that measured for the membrane-adsorbed αSyn where the C-termini were further appart. This result suggest that the interactions between C-termini are different when adsorbed onto a membrane than in a fibril. This could be either due to the higher mobility of proteins, or due to specific interactions between C-termini in lipid-adsorbed αSyn. If it was the latter, PICUP could hold the key to understanding the organization of αSyn when adsorbed to lipids, and shed light in the cooperatibity of its binding^30^. Given its sensitivity to proximity between monomers and their orientation, PICUP could also prove a valuable method to study differences in structure between fibril morphs.

### 4.4. Conclusion

We have showed that the cross-linking of αSyn varies depending on the dimensionality of the system studied (Figure 9). In solution (3D), a combination of the long-range interactions between the C and N termini of the monomer, and the interaction between C-termini of monomers in an oligomer, gives rise to a complex cross-linking pattern. When αSyn is adsorbed to lipids, in 2D, the internal cross-linking is blocked, and we observe transient interactions between C termini due to either chain diffusion of this mobile part or specific interaction between Ct tails. Finally, in fibrils, the system is reduced to 1D, in which monomers can only interact with their direct neighbours, giving rise to less cross-linking. These measurements reflect on αSyn’s behaviour as a monomer, as well as the transient interactions that occur in its self-and co-assembly. This study shows that PICUP is a very valuable tool that should be considered by anyone interested in the study of transient interactions of αSyn.

## References

1. Spillantini, M. G. et al. α-Synuclein in Lewy bodies. Nature 388, 839–840 (1997).

2. Polymeropoulos, M. H. et al. Mapping of a gene for Parkinson’s disease to chromosome 4q21-q23. Science (1979) 274, 1197–1199 (1996).

3. Pfefferkorn, C. M. et al. Depth of α-synuclein in a bilayer determined by fluorescence, neutron reflectometry, and computation. Biophys J 102, 613–621 (2012).

4. Hellstrand, E. et al. Adsorption of α-synuclein to supported lipid bilayers: Positioning and role of electrostatics. ACS Chem Neurosci 4, 1339–1351 (2013).

5. Hellstrand, E., Nowacka, A., Topgaard, D., Linse, S. & Sparr, E. Membrane Lipid Co-Aggregation with α-Synuclein Fibrils. PLoS One 8, (2013).

6. Buell, A. K. et al. Solution conditions determine the relative importance of nucleation and growth processes in α-synuclein aggregation. Proc Natl Acad Sci U S A 111, 7671–7676 (2014).

7. Galvagnion, C. et al. Lipid vesicles trigger α-synuclein aggregation by stimulating primary nucleation. Nat Chem Biol 11, 229–234 (2015).

8. Khare, S. D., Chinchilla, P. & Baum, J. Multifaceted interactions mediated by intrinsically disordered regions play key roles in alpha synuclein aggregation. Curr Opin Struct Biol 80, (2023).

9. Makasewicz, K., Linse, S. & Sparr, E. Interplay of α-synuclein with Lipid Membranes: Cooperative Adsorption, Membrane Remodeling and Coaggregation. JACS Au Preprint at 10.1021/jacsau.3c00579 (2023).

10. Siddiqui, I. J., Pervaiz, N. & Abbasi, A. A. The Parkinson Disease gene SNCA: Evolutionary and structural insights with pathological implication. Sci Rep 6, 24475 (2016).

11. Krüger, R. et al. AlaSOPro mutation in the gene encoding α-synuclein in Parkinson’s disease. Nat Genet 18, 106–108 (1998).

12. Zarranz, J. J. et al. The New Mutation, E46K, of α-Synuclein Causes Parkinson and Lewy Body Dementia. Ann Neurol 55, 164–173 (2004).

13. Appel-Cresswell, S. et al. Alpha-synuclein p.H50Q, a novel pathogenic mutation for Parkinson’s disease. Movement Disorders 28, 811–813 (2013).

14. Lesage, S. et al. G51D α-synuclein mutation causes a novel Parkinsonian-pyramidal syndrome. Ann Neurol 73, 459–471 (2013).

15. Pasanen, P. et al. A novel α-synuclein mutation A53E associated with atypical multiple system atrophy and Parkinson’s disease-type pathology. Neurobiol Aging 35, 2180.e1–2180.e5 (2014).

16. Yoshino, H. et al. Homozygous alpha-synuclein p.A53V in familial Parkinson’s disease. Neurobiol Aging 57, 248.e7–248.e12 (2017).

17. Newberry, R. W., Leong, J. T., Chow, E. D., Kampmann, M. & DeGrado, W. F. Deep mutational scanning reveals the structural basis for α-synuclein activity. Nat Chem Biol 16, 653–659 (2020).

18. Fusco, G. et al. Direct observation of the three regions in α-synuclein that determine its membrane-bound behaviour. Nat Commun 5, 3827 (2014).

19. Pálmadóttir, T., Malmendal, A., Leiding, T., Lund, M. & Linse, S. Charge Regulation during Amyloid Formation of α-Synuclein. J Am Chem Soc 143, 7777–7791 (2021).

20. Stephens, A. D. et al. Extent of N-terminus exposure of monomeric alpha-synuclein determines its aggregation propensity. Nat Commun 11, 2820 (2020).

21. Gaspar, R., Lund, M., Sparr, E. & Linse, S. Anomalous Salt Dependence Reveals an Interplay of Attractive and Repulsive Electrostatic Interactions in α-synuclein Fibril Formation. QRB Discov 1, (2020).

22. Yang, X., Wang, B., Hoop, C. L., Williams, J. K. & Baum, J. NMR unveils an N-terminal interaction interface on acetylated-α-synuclein monomers for recruitment to fibrils. Proceedings of the National Academy of Sciences 118, e2017452118 (2021).

23. Farzadfard, A., et al. The C-terminal tail of α-synuclein protects against aggregate replication but is critical for oligomerization. Commun Biol 5, (2022).

24. Pálmadóttir, T. et al. Morphology-Dependent Interactions between α-Synuclein Monomers and Fibrils. Int J Mol Sci 24, 5191 (2023).

25. Weinreb, P. H., Zhen, W., Poon, A. W., Conway, K. A. & Lansbury, P. T. NACP, a protein implicated in Alzheimer’s disease and learning, is natively unfolded. Biochemistry 35, 13709– 13715 (1996).

26. Theillet, F. X. et al. Structural disorder of monomeric α-synuclein persists in mammalian cells. Nature 530, 45–50 (2016).

27. Davidson, W. S., Jonas, A., Clayton, D. F. & George, J. M. Stabilization of α-Synuclein secondary structure upon binding to synthetic membranes. Journal of Biological Chemistry 273, 9443–9449 (1998).

28. Bodner, C. R., Dobson, C. M. & Bax, A. Multiple Tight Phospholipid-Binding Modes of α-Synuclein Revealed by Solution NMR Spectroscopy. J Mol Biol 390, 775–790 (2009).

29. Roeters, S. J. et al. Elevated concentrations cause upright alpha-synuclein conformation at lipid interfaces. Nat Commun 14, (2023).

30. Makasewicz, K. et al. Cooperativity of α-Synuclein Binding to Lipid Membranes. ACS Chem Neurosci 12, 2099–2109 (2021).

31. Shahmoradian, S. H. et al. Lewy pathology in Parkinson’s disease consists of crowded organelles and lipid membranes. Nat Neurosci 22, 1099–1109 (2019).

32. Heise, H. et al. Molecular-level secondary structure, polymorphism, and dynamics of full-length α-synuclein fibrils studied by solid-state NMR. Proceedings of the National Academy of Sciences 102, 15871–15876 (2005).

33. Li, B. et al. Cryo-EM of full-length α-synuclein reveals fibril polymorphs with a common structural kernel. Nat Commun 9, (2018).

34. Vilar, M. et al. The fold of α-synuclein fibrils. Proceedings of the National Academy of Sciences 105, (2008).

35. Gaspar, R. et al. Secondary nucleation of monomers on fibril surface dominates α-synuclein aggregation and provides autocatalytic amyloid amplification. Q Rev Biophys 50, (2017).

36. Cohen, S. I. A. et al. Distinct thermodynamic signatures of oligomer generation in the aggregation of the amyloid-β peptide. Nat Chem 10, 523–531 (2018).

37. Dear, A. J. et al. Kinetic diversity of amyloid oligomers. 117, 12087–12094 (2020).

38. Conway, K. A. et al. Acceleration of oligomerization, not fibrillization, is a shared property of both α-synuclein mutations linked to early-onset Parkinson’s disease: implications for pathogenesis and therapy. Proceedings of the National Academy of Sciences 97, 571–576 (2000).

39. Winner, B. et al. In vivo demonstration that α-synuclein oligomers are toxic. Proc Natl Acad Sci U S A 108, 4194–4199 (2011).

40. Walsh, D. M. et al. Naturally secreted oligomers of amyloid β protein potently inhibit hippocampal long-term potentiation in vivo. Nature 416, 535–539 (2002).

41. Kayed, R. et al. Common structure of soluble amyloid oligomers implies common mechanism of pathogenesis. Science (1979) 300, 486–489 (2003).

42. Andreasen, M., Lorenzen, N. & Otzen, D. Interactions between misfolded protein oligomers and membranes: A central topic in neurodegenerative diseases? Biochimica et Biophysica Acta - Biomembranes vol. 1848 1897–1907 Preprint at 10.1016/j.bbamem.2015.01.018 (2015).

43. Rodriguez Camargo, D. C., et al. Surface-Catalyzed Secondary Nucleation Dominates the Generation of Toxic IAPP Aggregates. Front Mol Biosci 8, 1037 (2021).

44. Campioni, S. et al. A causative link between the structure of aberrant protein oligomers and their toxicity. Nat Chem Biol 6, 140–147 (2010).

45. Dear, A. J. et al. Identification of on- and off-pathway oligomers in amyloid fibril formation. Chem Sci 11, 6236–6247 (2020).

46. Arosio, P., Knowles, T. P. J. & Linse, S. On the lag phase in amyloid fibril formation. Physical Chemistry Chemical Physics 17, 7606–7618 (2015).

47. Cawood, E. E., Karamanos, T. K., Wilson, A. J. & Radford, S. E. Visualizing and trapping transient oligomers in amyloid assembly pathways. Biophys Chem 268, (2021).

48. Fancy, D. A. & Kodadek, T. Chemistry for the analysis of protein-protein interactions: Rapid and efficient cross-linking triggered by long wavelength light. Proceedings of the National Academy of Sciences 96, 6020–6024 (1999).

49. Li, Y. et al. Dissecting the role of disulfide bonds on the amyloid formation of insulin. Biochem Biophys Res Commun 423, 373–378 (2012).

50. Shahpasand-Kroner, H. et al. Three-repeat and four-repeat tau isoforms form different oligomers. Protein Science 31, 613–627 (2022).

51. Bitan, G., Lomakin, A. & Teplow, D. B. Amyloid β-protein oligomerization: Prenucleation interactions revealed by photo-induced cross-linking of unmodified proteins. Journal of Biological Chemistry 276, 35176–35184 (2001).

52. Fancy, D. A. et al. Scope, limitations and mechanistic aspects of the photo-induced cross-linking of proteins by water-soluble metal complexes. Chem Biol 7, 697–708 (2000).

53. Bitan, G. Structural Study of Metastable Amyloidogenic Protein Oligomers by Photo-Induced Cross-Linking of Unmodified Proteins. Methods Enzymol 413, 217–236 (2006).

54. Li, H.-T., Lin, X.-J., Xie, Y.-Y. & Hu, H.-Y. The Early Events of α-Synuclein Oligomerization Revealed by Photo-Induced Cross-Linking. Protein Pept Lett 13, 385–390 (2006).

55. Ortigosa-Pascual, L., Leiding, T., Linse, S. & Pálmadóttir, T. Photo-Induced Cross-Linking of Unmodified α-Synuclein Oligomers. ACS Chem Neurosci 14, (2023).

56. Borsarelli, C. D. et al. Biophysical properties and cellular toxicity of covalent crosslinked oligomers of α-synuclein formed by photoinduced side-chain tyrosyl radicals. Free Radic Biol Med 53, 1004–1015 (2012).

57. Roth, M. Fluorescence reaction for amino acids. Anal Chem 43, 880–882 (1971).

58. Benson, J. R. & Hare, P. E. O-phthalaldehyde: fluorogenic detection of primary amines in the picomole range. Comparison with fluorescamine and ninhydrin. Proceedings of the National Academy of Sciences 72, 619–622 (1975).

59. Axell, E. et al. The role of shear forces in primary and secondary nucleation of amyloid fibrils. Proceedings of the National Academy of Sciences 121, (2024).

60. Curry, A. M. et al. Mapping of photochemically-derived dityrosine across fe-bound n-acetylated α-synuclein. Life 10, 1–12 (2020).

61. Cohen, S. I. A., Vendruscolo, M., Dobson, C. M. & Knowles, T. P. J. From macroscopic measurements to microscopic mechanisms of protein aggregation. Journal of Molecular Biology vol. 421 160–171 Preprint at 10.1016/j.jmb.2012.02.031 (2012).

62. Jo, E., McLaurin, J. A., Yip, C. M., St. George-Hyslop, P. & Fraser, P. E. α-Synuclein membrane interactions and lipid specificity. Journal of Biological Chemistry 275, 34328– 34334 (2000).

63. Eliezer, D., Kutluay, E., Bussell, R. & Browne, G. Conformational properties of α-synuclein in its free and lipid-associated states. J Mol Biol 307, 1061–1073 (2001).

64. Skaff, O., Jolliffe, K. A. & Hutton, C. A. Synthesis of the side chain cross-linked tyrosine oligomers dityrosine, trityrosine, and pulcherosine. Journal of Organic Chemistry 70, 7353– 7363 (2005).

65. Dedmon, M. M., Lindorff-Larsen, K., Christodoulou, J., Vendruscolo, M. & Dobson, C. M. Mapping long-range interactions in α-synuclein using spin-label NMR and ensemble molecular dynamics simulations. J Am Chem Soc 127, 476–477 (2005).

66. Zhang, C. et al. C-terminal truncation modulates α-Synuclein’s cytotoxicity and aggregation by promoting the interactions with membrane and chaperone. Commun Biol 5, 798 (2022).

67. Bertoncini, C. W. et al. Release of long-range tertiary interactions potentiates aggregation of natively unstructured α-synuclein. 102, 1430–1435 (2005).

68. Cho, M. K. et al. Structural characterization of α-synuclein in an aggregation prone state. Protein Science 18, 1840–1846 (2009).

69. Zhou, W. et al. Methionine oxidation stabilizes non-toxic oligomers of α-synuclein through strengthening the auto-inhibitory intra-molecular long-range interactions. Biochim Biophys Acta Mol Basis Dis 1802, 322–330 (2010).

70. Antony, T. et al. Cellular polyamines promote the aggregation of α-synuclein. Journal of Biological Chemistry 278, 3235–3240 (2003).

71. Fernández, C. O. et al. NMR of α-synuclein-polyamine complexes elucidates the mechanism and kinetics of induced aggregation. EMBO Journal 23, 2039–2046 (2004).

72. Ramis, R., Ortega-Castro, J., Vilanova, B., Adrover, M. & Frau, J. Cu2+, Ca2+, and methionine oxidation expose the hydrophobic α-synuclein NAC domain. Int J Biol Macromol 169, 251–263 (2021).

73. Baba, M. et al. Aggregation of alpha-synuclein in Lewy bodies of sporadic Parkinson’s disease and dementia with Lewy bodies. Am J Pathol 152, 879 (1998).

74. Li, W. et al. Aggregation promoting C-terminal truncation of α-synuclein is a normal cellular process and is enhanced by the familial Parkinson’s disease-linked mutations. Proceedings of the National Academy of Sciences 102, 2162–2167 (2005).

75. Muntané, G., Ferrer, I. & Martinez-Vicente, M. α-synuclein phosphorylation and truncation are normal events in the adult human brain. Neuroscience 200, 106–119 (2012).

76. Bhattacharjee, P. et al. Mass Spectrometric Analysis of Lewy Body-Enriched α-Synuclein in Parkinson’s Disease. J Proteome Res 18, 2109–2120 (2019).

77. Crowther, R. A., Jakes, R., Spillantini, M. G. & Goedert, M. Synthetic filaments assembled from C-terminally truncated α-synuclein. FEBS Lett 436, 309–312 (1998).

78. Murray, I. V. J. et al. Role of α-synuclein carboxy-terminus on fibril formation in vitro. Biochemistry 42, 8530–8540 (2003).

79. Levitan, K. et al. Conserved C-terminal charge exerts a profound influence on the aggregation rate of α-synuclein. J Mol Biol 411, 329–333 (2011).

80. Van Der Wateren, I. M., Knowles, T. P. J., Buell, A. K., Dobson, C. M. & Galvagnion, C. C-terminal truncation of α-synuclein promotes amyloid fibril amplification at physiological pH. Chem Sci 9, 5506–5516 (2018).

81. Ni, X., McGlinchey, R. P., Jiang, J. & Lee, J. C. Structural Insights into α-Synuclein Fibril Polymorphism: Effects of Parkinson’s Disease-Related C-Terminal Truncations. J Mol Biol 431, 3913–3919 (2019).

82. McGlinchey, R. P. et al. C-terminal α-synuclein truncations are linked to cysteine cathepsin activity in Parkinson’s disease. Journal of Biological Chemistry 294, 9973–9984 (2019).

83. Kanda, S., Bishop, J., Eglitis, M., Yang, Y. & Mouradian, M. Enhanced vulnerability to oxidative stress by α-synuclein mutations and C-terminal truncation. Neuroscience 97, 279– 284 (2000).

84. Stefanova, N., Klimaschewski, L., Poewe, W., Wenning, G. K. & Reindl, M. Glial cell death induced by overexpression of α-synuclein. J Neurosci Res 65, 432–438 (2001).

85. Ma, L. et al. C-terminal truncation exacerbates the aggregation and cytotoxicity of α-Synuclein: A vicious cycle in Parkinson’s disease. Biochim Biophys Acta Mol Basis Dis 1864, 3714–3725 (2018).

86. Yagi-Utsumi, M., Satoh, T. & Kato, K. Structural basis of redox-dependent substrate binding of protein disulfide isomerase. Sci Rep 5, 13909 (2015).

87. Kumari, P. et al. Structural insights into α-synuclein monomer-fibril interactions. Proceedings of the National Academy of Sciences 118, (2021).

88. Sahin, C. et al. Structural Basis for Dityrosine-Mediated Inhibition of α-Synuclein Fibrillization. J Am Chem Soc 144, 11949–11954 (2022).

89. Apetri, M. M., Maiti, N. C., Zagorski, M. G., Carey, P. R. & Anderson, V. E. Secondary structure of α-synuclein oligomers: Characterization by Raman and atomic force microscopy. J Mol Biol 355, 63–71 (2006).

90. Cremades, N. et al. Direct observation of the interconversion of normal and toxic forms of α-synuclein. Cell 149, 1048–1059 (2012).

91. Celej, M. S. et al. Toxic prefibrillar α-synuclein amyloid oligomers adopt a distinctive antiparallel β-sheet structure. Biochemical Journal 443, 719–726 (2012).

92. Lorenzen, N. et al. The role of stable α-synuclein oligomers in the molecular events underlying amyloid formation. J Am Chem Soc 136, 3859–3868 (2014).

93. Chen, S. W. et al. Structural characterization of toxic oligomers that are kinetically trapped during α-synuclein fibril formation. Proc Natl Acad Sci U S A 112, E1994–E2003 (2015).

94. Fusco, G. et al. Structural basis of membrane disruption and cellular toxicity by α-synuclein oligomers. Science (1979) 358, 1440–1443 (2017).

95. Ehrnhoefer, D. E. et al. EGCG redirects amyloidogenic polypeptides into unstructured, off-pathway oligomers. Nat Struct Mol Biol 15, 558–566 (2008).

96. Fusco, G., Sanz-Hernandez, M. & De Simone, A. Order and disorder in the physiological membrane binding of α-synuclein. Curr Opin Struct Biol 48, 49–57 (2018).

97. Petrache, H. I. et al. Structure and Fluctuations of Charged Phosphatidylserine Bilayers in the Absence of Salt. Biophys J 86, 1574–1586 (2004).

98. Nagle, J. F. & Tristram-Nagle, S. Structure of lipid bilayers. Biochimica et Biophysica Acta (BBA) - Reviews on Biomembranes 1469, 159–195 (2000).

99. Corey, R. B. & Pauling, L. C. Fundamental dimensions of polypeptide chains. Proceedings of the Royal Society of London. Series B-Biological Sciences 141, 10–20 (1953).

100. Kohn, J. E. et al. Random-coil behavior and the dimensions of chemically unfolded proteins. Proceedings of the National Academy of Sciences 101, 12491–12496 (2004).

101. Riek, R. & Eisenberg, D. S. The activities of amyloids from a structural perspective. Nature 539, 227–235 (2016).

102. Tuttle, M. D. et al. Solid-state NMR structure of a pathogenic fibril of full-length human α-synuclein. Nat Struct Mol Biol 23, 409–415 (2016).

