## Supporting Information for "On the transient interactions of α-synuclein in different dimensions"

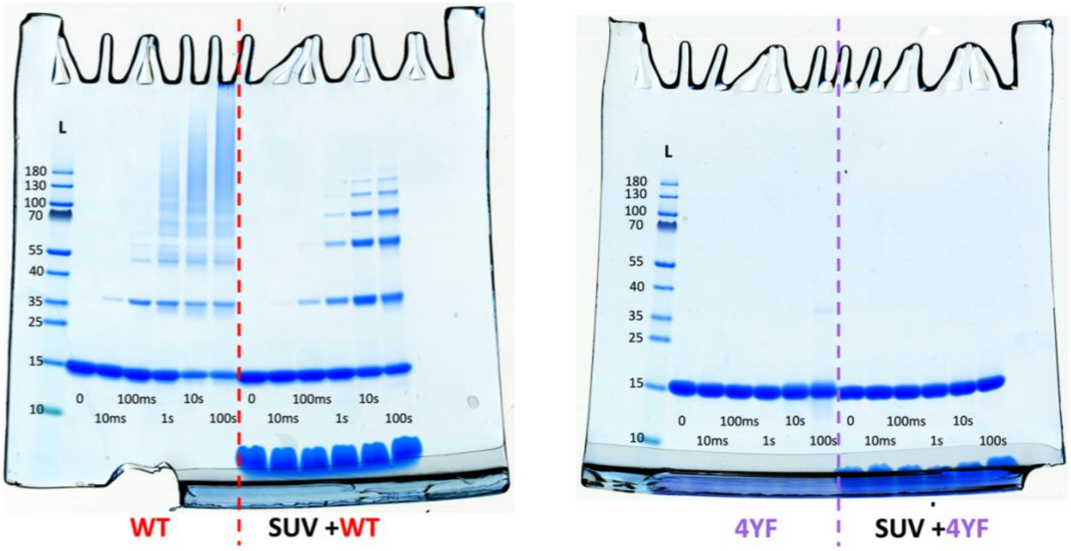


**Figure S1**. Full gels of pictures shown in Figure 2A, 2B and 5B. Cross-linking of WT and 4YF variant of αSyn, either without (left) or with (right) SUVs added, at a L/P ratio of 250. Lighting time is indicated in numbers below each lane.

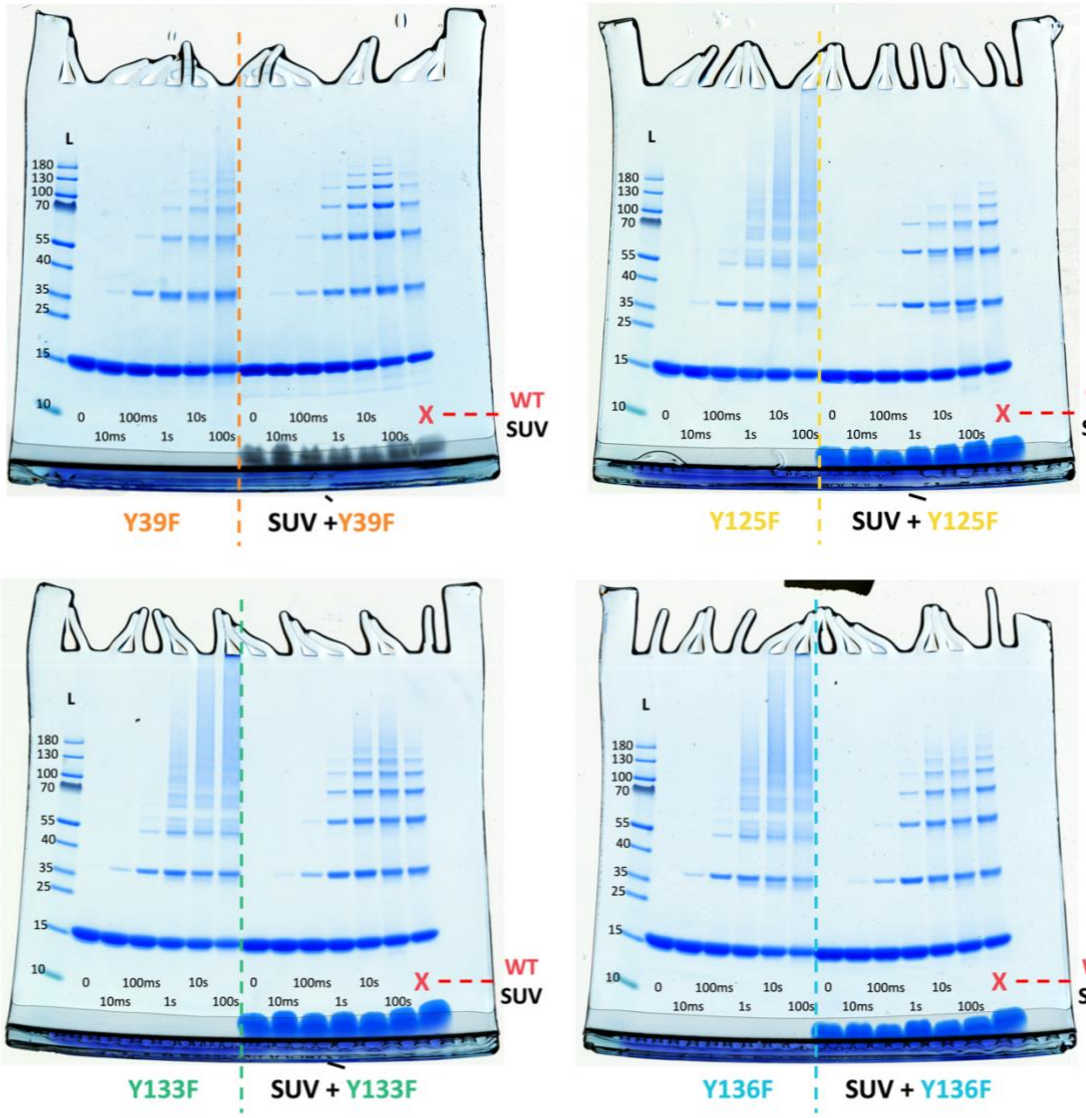


**Figure S2**. Full gels of pictures shown in Figure 3 and 6. Cross-linking of αSyn mutants Y39F, Y125F, Y133F and Y136F, either without (left) or with (right) SUVs added, at a L/P ratio of 250. Lighting time is indicated in numbers below each lane. All gels contain a reference of WT αSyn sample in presence o SUV (L/P = 250), cross-linked for 10 s.

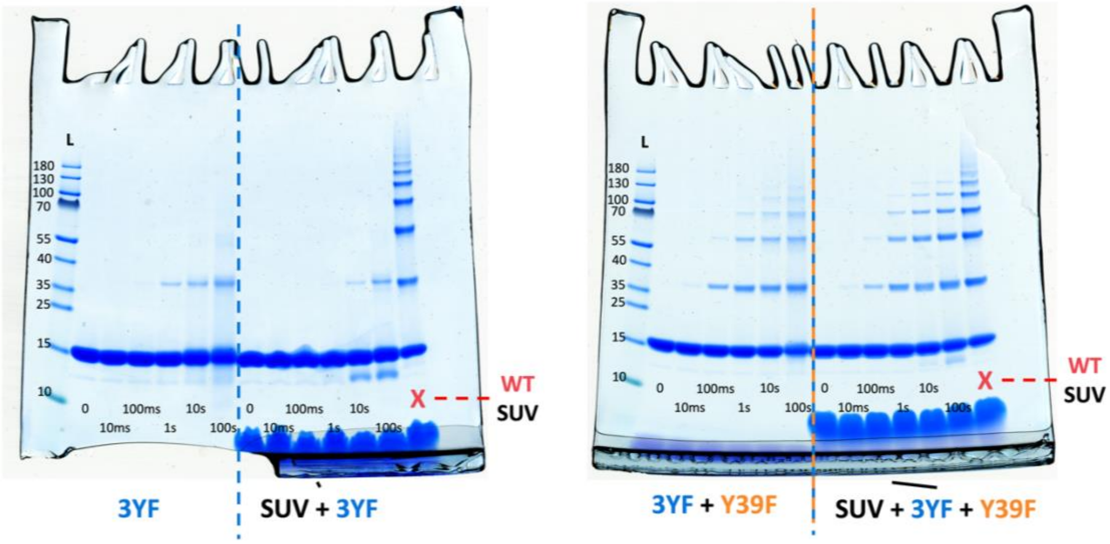


**Figure S3**. Full gels of pictures shown in Figure 4. Cross-linking of αSyn mutants 3YF, and a 50:50 ratio mixture of 3YF and Y39F, either without (left) or with (right) SUVs added, at a L/P ratio of 250. Lighting time is indicated in numbers below each lane. All gels contain a reference of WT αSyn sample in presence o SUV (L/P = 250), cross-linked for 10 s.

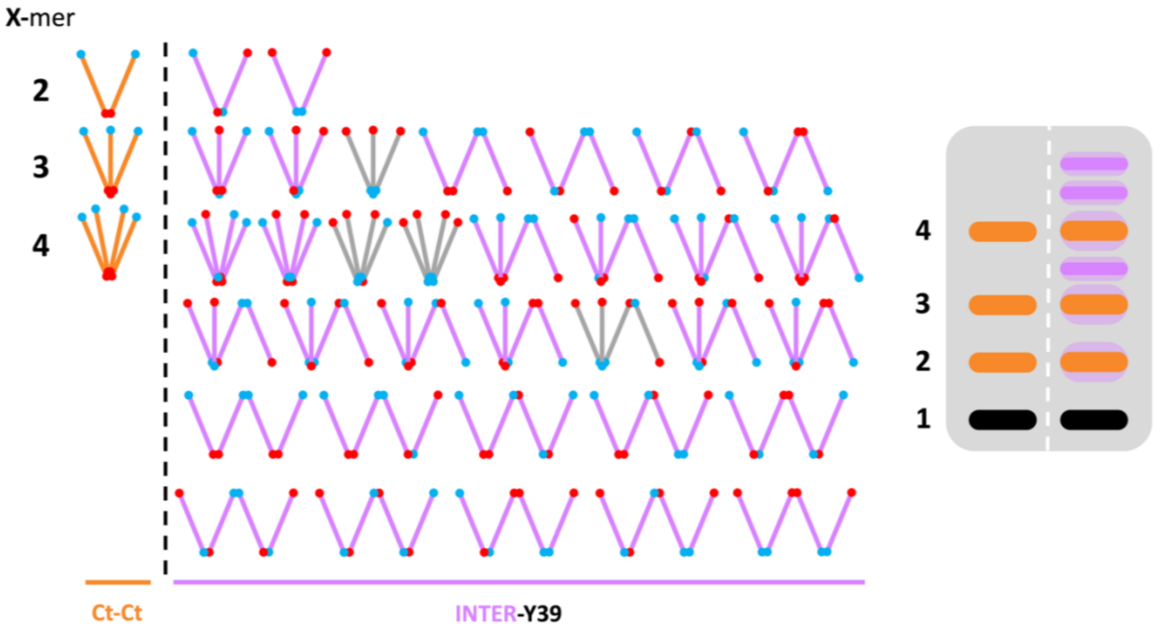


**Figure S4**. The effect of addition of Y39 in the complexity of cross-linked morphology by contribution to inter-molecular cross-linking. This figure represents the same as Figure 8A, but considering the Nt and Ct as different. Due to that, the complexity of obtained species highly increases. This leads to species with similar morphology, yet different cross-linking, and therefore,potentially slightly different migration (i.e. the three morphologies for dimer are the same, but since the Nt and Ct are slightly different distance from their respective ends of the peptide, they migh migrate differently). This is represented by a faded purple background around the orange bands. This is most likely closer to the real system than that represented in Figure 8A, as we see the shadow of a dimer band in solution, which disappears with increasing L/P, as mentioned in discussion. One should note this increased complexity wen considering the termini separately also applies to the system where Y39 contributes to intra-molecular cross-linking (Figure 8B).
